# Plasma membrane nano-organization specifies phosphoinositide effects on Rho-GTPases and actin dynamics in tobacco pollen tubes

**DOI:** 10.1101/2020.08.12.248419

**Authors:** Marta Fratini, Praveen Krishnamoorthy, Irene Stenzel, Mara Riechmann, Kirsten Bacia, Mareike Heilmann, Ingo Heilmann

## Abstract

Pollen tube growth requires coordination of cytoskeletal dynamics and apical secretion. The regulatory phospholipid, phosphatidylinositol 4,5-bisphosphate (PtdIns(4,5)P_2_) is enriched in the subapical plasma membrane of pollen tubes and can influence both actin dynamics and secretion. How alternative PtdIns(4,5)P_2_-effects are specified is unclear. Spinning disc microscopy (SD) reveals dual distribution of a fluorescent PtdIns(4,5)P_2_-reporter in dynamic plasma membrane nanodomains vs. apparent diffuse membrane labelling, consistent with spatially distinct coexisting pools of PtdIns(4,5)P_2_. Several PI4P 5-kinases (PIP5Ks) can generate PtdIns(4,5)P_2_ in pollen tubes. Despite localizing to one membrane region, AtPIP5K2 and NtPIP5K6 display distinctive overexpression effects on cell morphologies, respectively related to altered actin dynamics or membrane trafficking. When analyzed by SD, AtPIP5K2-EYFP associated with nanodomains, whereas NtPIP5K6-EYFP localized diffusely. Chimeric AtPIP5K2 and NtPIP5K6 variants with reciprocally swapped membrane-associating domains evoked reciprocally shifted effects on cell morphology upon overexpression. Overall, PI4P 5-kinase variants targeted to nanodomains stabilized actin, suggesting a specific function of PtdIns(4,5)P_2_-nanodomains. A distinct role of nanodomain-associated AtPIP5K2 in actin regulation is further supported by proximity to and interaction with the Rho-GTPase NtRac5, and by functional interplay with elements of ROP-signalling. Plasma membrane nano-organization may thus aid the specification of PtdIns(4,5)P_2_-functions to coordinate cytoskeletal dynamics and secretion in pollen tubes.

## Introduction

The extreme cell shapes of plant pollen tubes or root hairs arise from polar tip growth (Hepler and Winship, 2015; Orr et al., 2020). For polar tip growth, polarized cell expansion is achieved by the tip-focused transport of vesicles containing cell wall material and membrane towards the expanding cell apex, their secretion at a narrow area of the cell surface and - upon cargo delivery - the endocytotic retrieval of a majority of vesicles for recycling (Cheung and Wu, 2008). The balance of apical secretion and recycling of vesicles requires the coordination of membrane trafficking and cytoskeletal dynamics at the plasma membrane (Grebnev et al., 2017; Guo and Yang, 2020; Orr et al., 2020).

The plasma membrane serves multiple physiological functions, including as a hydrophobic barrier, for selective transport, signal transduction, attachment of the cytoskeleton and for membrane trafficking. Its body is composed of amphiphilic lipids, including glycerophospholipids, sphingolipids and sterols (Furt et al., 2011; Mamode Cassim et al., 2019). Some lipid classes are highly abundant, such as the structural glycerophospholipids phosphatidylcholine or phosphatidylethanolamine, whereas other lipid classes occur in much smaller proportions, such as phosphatidylserine or other anionic lipids, such as phosphoinositides (Furt et al., 2011), which are in the focus of this study. Phosphoinositides are a minor class of regulatory phospholipids that derive from phosphatidylinositol by the phosphorylation of the inositol head group and in eukaryotes contribute to the control of membrane trafficking and cytoskeletal dynamics (Heilmann and Heilmann, 2015). Importantly, phosphoinositides can bind different protein partners at the cytosolic face of the plasma membrane (Heilmann, 2016; Gerth et al., 2017; Noack and Jaillais, 2020), thereby influencing alternative processes. It is an important unresolved question of eukaryotic cell biology how phosphoinositides exert specific regulatory effects.

A role of phosphoinositides in the control of polar tip growth has been reported by numerous studies on root hairs (Vincent et al., 2005; Kusano et al., 2008; Stenzel et al., 2008; Stanislas et al., 2015) or pollen tubes (Kost et al., 1999; Monteiro et al., 2005b; Monteiro et al., 2005a; Ischebeck et al., 2008; Kost, 2008; Sousa et al., 2008; Ischebeck et al., 2010a; Ischebeck et al., 2010b; Ischebeck et al., 2011; Stenzel et al., 2012; Heilmann and Heilmann, 2015; Bloch et al., 2016); or moss filaments (Saavedra et al., 2011; Saavedra et al., 2015); or even fungal hyphae (Ischebeck et al., 2010a; Mähs et al., 2012). While phosphoinositides evidently contribute to the control and coordination of various aspects of polar tip growth in plants, the factors determining their individual specific effects have remained elusive.

The subcellular distribution of some phosphoinositides has been visualized in plant cells by genetically encoded fluorescent reporters, for instance such based on the Pleckstrin homology (PH)-domain of mammalian phospholipase C (PLC)δ1 specifically recognizing phosphatidylinositol 4,5-bisphosphate (PtdIns(4,5)P_2_) (Varnai and Balla, 1998; Kost et al., 1999; Balla and Varnai, 2002; van Leeuwen et al., 2007; Simon et al., 2014) or other lipid binding modules specific for phosphatidylinositol 4-phosphate (PtdIns4P) (Vermeer et al., 2009; Simon et al., 2014; Simon et al., 2016). In pollen tubes, fluorescent biosensors show PtdIns(4,5)P_2_ enriched in a subapical plasma membrane region (Kost et al., 1999), where the lipid has been proposed to influence both membrane trafficking and cytoskeletal dynamics (Malho et al., 2006; Kost, 2008; Ischebeck et al., 2010a; Heilmann and Heilmann, 2015; Gerth et al., 2017). The PtdIns(4,5)P_2_-rich region of the subapical plasma membrane spans from approx. 1-13 µm as measured from the tip of growing tobacco pollen tubes, spanning a region accumulating secretory and recycling vesicles that is characterized by a fine-tuned actin network required for proper trafficking (Hepler and Winship, 2015; Heilmann and Ischebeck, 2016; Grebnev et al., 2017).

PtdIns(4,5)P_2_ in this region of the pollen tube plasma membrane is essential for polar tip-growth, because genetic lesions in PtdIns(4,5)P_2_ biosynthesis result in massively decreased pollen germination and in impaired pollen tube expansion (Ischebeck et al., 2008; Sousa et al., 2008; Heilmann and Ischebeck, 2016). As loss of function mutants show defects in pollen germination, regulatory effects of PtdIns(4,5)P_2_ have been assessed mostly by gain of function approaches, in which the perturbation of PtdIns(4,5)P_2_ biosynthesis also results in defective polar tip growth. At least two categories of characteristic aberrant cell morphologies arise from the overexpression of different PI4P 5-kinases in pollen tubes. For instance, the overexpression of the Arabidopsis PI4P 5-kinases AtPIP5K4 (Ischebeck et al., 2008; Sousa et al., 2008; Ischebeck et al., 2010b), AtPIP5K5 (Ischebeck et al., 2008; Ischebeck et al., 2010b), AtPIP5K6 (Zhao et al., 2010), or tobacco NtPIP5K6 (Stenzel et al., 2012) results in a characteristic and irregular tip branching phenotype of the pollen tubes, which is accompanied by increased apical deposition of pectin. The associated cell morphologies have previously been summarized as “secretion phenotypes” (Ischebeck et al., 2010b). By contrast, these effects are not apparent when other PI4P 5-kinases are overexpressed, such as AtPIP5K2 (Stenzel et al., 2012), AtPIP5K10 (Ischebeck et al., 2011) or AtPIP5K11 (Ischebeck et al., 2011), which instead cause extensive pollen tube tip swelling upon overexpression, proposed to be linked to changes in actin (Ischebeck et al., 2011). It is important to note that no increased pectin deposition or tip branching is observed upon overexpression of AtPIP5K2, AtPIP5K10 and AtPIP5K11 (Ischebeck et al., 2011; Stenzel et al., 2012), suggesting that these enzymes are functionally divergent (Stenzel et al., 2012).

The differences in reported effects of PI4P 5-kinases on cell morphology make the pollen tube a unique model to study the determinants for the specificity of PtdIns(4,5)P_2_ effects within a well-defined plasma membrane region of one cell (Ischebeck et al., 2010a; Stenzel et al., 2012; Heilmann and Heilmann, 2015; Gerth et al., 2017). As catalytic activity of PI4P 5-kinases was required to mediate the respective effects on cell morphologies (Ischebeck et al., 2008; Ischebeck et al., 2011), we have previously proposed the presence of at least two functional pools of PtdIns(4,5)P_2_ within the subapical plasma membrane of pollen tube cells (Ischebeck et al., 2011; Stenzel et al., 2012; Heilmann and Heilmann, 2013, 2015; Heilmann, 2016; Gerth et al., 2017), which may be defined by the respective PI4P 5-kinases mediating their biosynthesis. Such PtdIns(4,5)P_2_ pools may preferentially influence membrane trafficking or cytoskeletal dynamics, respectively (Stenzel et al., 2012). However, direct experimental evidence for or against the pool hypothesis has been elusive so far, and no systematic study has been conducted to compare pollen PI4P 5-kinases to address their proposed distinct functionalities side-by-side.

In *Arabidopsis thaliana*, PI4P 5-kinases are represented as an enzyme family of 11 members (Supplemental Fig. S1), which can be divided into subfamilies A (PIP5K10 and PIP5K11) and B (PIP5K1-PIP5K9), based on differences in protein architecture. All isoforms preferentially convert the substrate phosphatidylinositol 4-phosphate (PtdIns4P) to PtdIns(4,5)P_2_ in vitro (Ischebeck et al., 2008; Stenzel et al., 2008; Ischebeck et al., 2011; Stenzel et al., 2012; Ischebeck et al., 2013). Enzymes of subfamily A lack several N-terminal domains found in their counterparts of subfamily B (Fig. 1A) and have so far only been found in Arabidopsis. PI4P 5-kinases of subfamily A are only weakly expressed and Arabidopsis *pip5k10 pip5k11* double mutants displayed only minor defects in pollen tube growth (Ischebeck et al., 2011). Overall, the physiological relevance of the “small” PI4P 5-kinases of subfamily A remains thus unclear. By contrast, Arabidopsis mutants with defects in genes for PI4P 5-kinases of subfamily B display massively reduced pollen germination and reduced pollen tube growth, such as reported for the Arabidopsis *pip5k4 pip5k5* mutants (Ischebeck et al., 2008; Sousa et al., 2008), highlighting a more pronounced contribution to pollen tube growth. Importantly, pollen-expressed members of the PI4P 5-kinase subfamily B have previously been found to mediate distinct effects either on membrane trafficking, such as AtPIP5K4 and AtPIP5K5 (Ischebeck et al., 2008; Sousa et al., 2008), AtPIP5K6 (Zhao et al., 2010) or NtPIP5K6 (Stenzel et al., 2012), or on the cytoskeleton, such as AtPIP5K2 (Stenzel et al., 2012), raising the question how PI4P 5-kinases of subfamily B exert isoform-specific regulatory effects.

**Figure 1.**
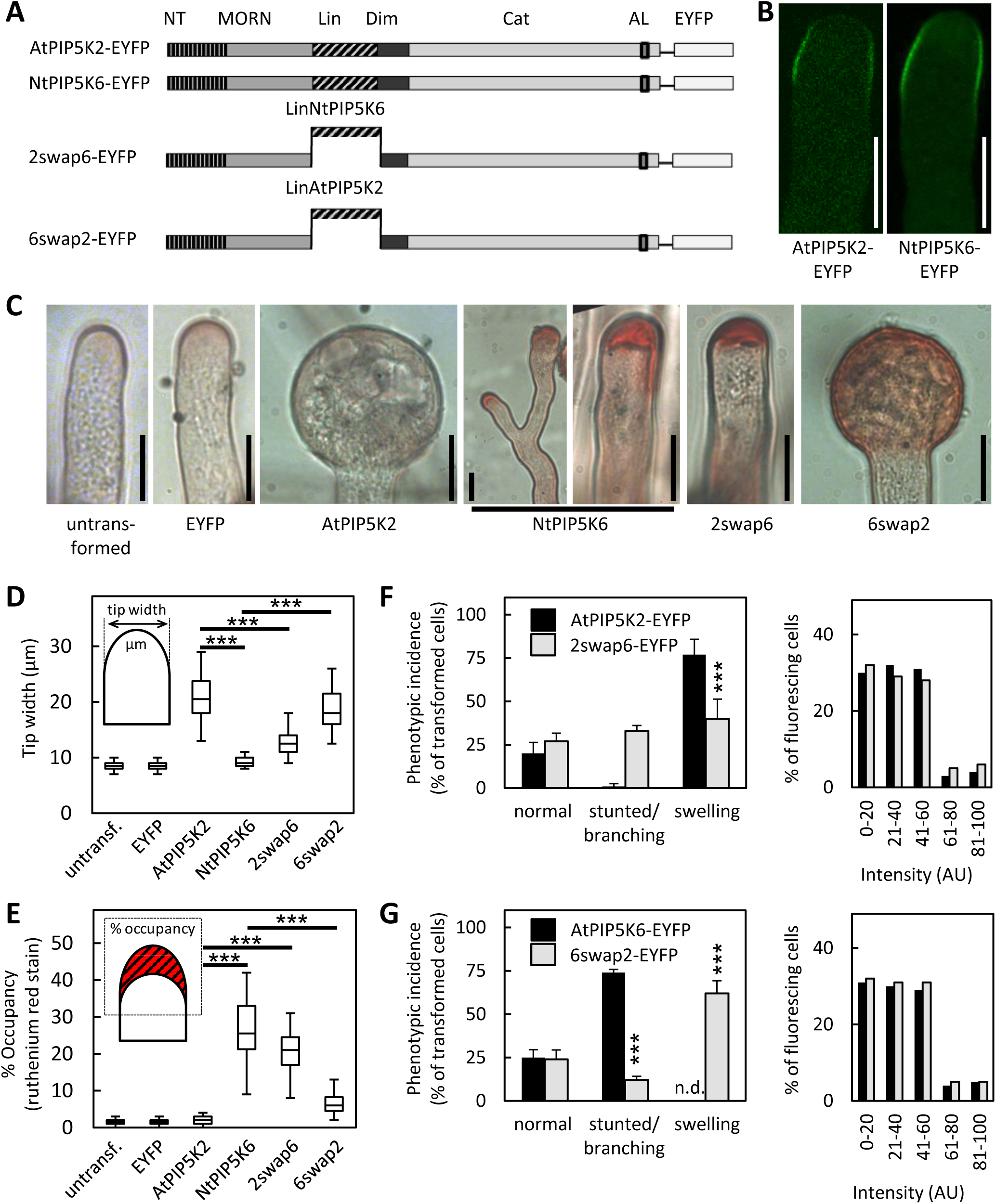
The PI4P 5-kinases, AtPIP5K2 and NtPIP5K6, have divergent regulatory effects in tobacco pollen tubes. **A**, AtPIP5K2 and NtPIP5K6 display the characteristic domain structure of PI4P 5-kinases of subfamily B. NT, N-terminal domain; MORN, membrane occupation and recognition nexus-repeat domain; Lin, linker domain; Dim, dimerization domain; Cat, catalytic domain; AL, activation loop. The chimeric enzymes 2swap6 and 6swap2 were created by reciprocally swapping the Lin-domain between AtPIP5K2 and NtPIP5K6, as indicated. EYFP, yellow fluorescent protein. **B**, AtPIP5K2-EYFP or NtPIP5K6-EYFP were transiently expressed in tobacco pollen tubes under the control of the Lat52-promoter, and the fluorescence distribution was recorded by confocal microscopy. Representative images show both enzymes localized to a subapical region of the plasma membrane of growing pollen tubes. Scale bar, 10 µm. **C**, The effects of overexpressing AtPIP5K2-EYFP, NtPIP5K6 or the chimeric variants 2swap6-EYFP or 6swap2-EYFP on the morphology of tobacco pollen tubes was assessed upon transient expression under the control of the Lat52 promoter. Representative cell morphologies are shown in comparison to untransformed cells or to pollen tubes expressing an EYFP control, as indicated, and include pollen tube tip swelling or the increased apical deposition of pectin, as stained by ruthenium red. Please note that all pollen tubes shown were stained by ruthenium red. Pollen tubes displaying increased apical pectin deposition also showed stunted or branched growth (not shown). Scale bar, 10 µm. **D, E**, The degree of pollen tube tip swelling (D) or of ruthenium red staining (E) were quantified by measuring tip with (n≥200) or the apical dye occupancy (n≥200), respectively, as indicated. **F, G**, The incidence of stunted growth/branching or tip swelling was recorded upon overexpression of AtPIP5K2-EYFP or 2swap6-EYFP (F), or upon overexpression of NtPIP5K6-EYFP or 6swap2-EYFP (G), respectively (n≥200 for all constructs analyzed). Right panels, Fluorescence intensity distribution corresponding to overexpression of the parental (black bars) or chimeric (grey bars) enzymes. The expression of all fusions was driven by the Lat52 promoter. Data in are given as means ± standard deviation. Asterists indicate a significant difference to the control, as indicated, according to a Student’s T-test. ***, p≤0.001.

PI4P 5-kinases of subfamily B share a unique domain structure consisting of an N-terminal (NT) domain, a membrane occupation and recognition nexus (MORN) repeat domain, a hypervariable linker (Lin)-domain and a catalytic (Cat) domain, which holds a variable insert region (Fig. 1A). This domain structure is only found in PI4P 5-kinases of plant origin and not in enzymes from non-plant organisms (Mueller-Roeber and Pical, 2002; Gerth et al., 2017). The well-defined domain architecture of PI4P 5-kinases of subfamily B enables the comparative analysis of features mediating isoform-specific functionality of the enzymes. To this end, previous work has focused on the comparison of the enzymes AtPIP5K2 and NtPIP5K6, which upon everexpression display pronounced functional differences and mutually exclusive effects on pollen tube morphologies: AtPIP5K2 causes pollen tube tip swelling and NtPIP5K6 evokes pronounced “secretion phenotypes” (Stenzel et al., 2012). To rationalize the functional divergence of AtPIP5K2 and NtPIP5K6, previous analyses have focused on the hypervariable Lin-domains, which contribute to the plasma membrane association of PI4P 5-kinases (Stenzel et al., 2012). Lin-domains are not conserved in sequence between PI4P 5-kinase isoforms, making them ideal candidate regions to mediate isoform-specific membrane recruitment (Stenzel et al., 2012; Heilmann and Heilmann, 2013; Gerth et al., 2017). The functionality of the Lin-domains is not understood in mechanistic detail, and the regions are predicted to be intrinsically disordered. Among PI4P 5-kinases of subfamily B, the functional distinction of AtPIP5K2 has previously been attributed to an increased accumulation of point mutations within the coding region for its Lin-domain, suggesting that the divergent evolution of membrane attachment regions might contribute to the rise of novel functionalities (Stenzel et al., 2012). When Lin-domains were previously swapped between AtPIP5K2 and NtPIP5K6, the exchange resulted in the reciprocal shift in the respectively arising cell morphologies (Stenzel et al., 2012). The combined observations suggest that one reason for the alternative physiological effects of the enzymes was recruitment by their Lin-domains into alternative regulatory contexts at the plasma membrane (Stenzel et al., 2012; Heilmann and Heilmann, 2013).

Different functional aspects of the plasma membrane are defined by the presence of certain protein complexes, resulting in a dynamic landscape of areas with regionally defined functional specifications (Jaillais and Ott, 2020). It is our working hypothesis that the recruitment of PI4P 5-kinases into one or another protein complex at the plasma membrane underlies the functional specificity of PtdIns(4,5)P_2_ formed by a given enzyme. Reciprocally, the localized formation of PtdIns(4,5)P_2_ may underlie the recruitment of proteins constituting a certain functional environment, possibly resulting in the self-organization of membrane nanodomains with specific functions. Numerous membrane-associated protein complexes mediate their function not diffusely in the plane of the membrane, but are focused in narrow spots (nanodomains, foci) of < 1 µm in diameter, which can be observed by spinning disc microscopy (SD). Examples include the assembly of the EXOCYST complex during secretion (Li et al., 2013; Zhang et al., 2013; Kalmbach et al., 2017; Sekereš et al., 2017) or the recruitment of adaptor proteins and the assembly of the clathrin coat during endocytosis (Konopka et al., 2008; Gadeyne et al., 2014; Bashline et al., 2015; Johnson and Vert, 2017). Similarly, monomeric GTPases of the Rho of plants (ROP) family, which are involved in the coordination of the actin cytoskeleton with secretion and endocytosis (Kost et al., 1999; Kost, 2008; Lee et al., 2008; Yalovsky et al., 2008; Grebnev et al., 2017), distribute into plasma membrane nanodomains (Platre et al., 2019). Interestingly, overexpression of the tobacco ROP, NtRac5, results in actin stabilization and pollen tube tip swelling (Klahre et al., 2006), suggesting a link between phosphoinositides and actin dynamics involving ROPs.

Despite a wealth of information on protein localization patterns and on protein-protein interactions within relevant protein complexes, the contribution of membrane lipids to the formation of particular membrane nanodomains has only recently begun to be unravelled (Platre et al., 2019). So far, there is only limited information on the lateral distribution of phospholipids within the plane of the plasma membrane in plants. In petunia pollen tubes, a fluorescence-tagged oxysterol binding protein decorated a spot-like pattern at the apical plasma membrane (Skirpan et al., 2006). Based on circumstantial evidence, the presence of PtdIns(4,5)P_2_-containing nanodomains of ∼70 nm in diameter has previously been suggested for plasma membrane vesicles isolated from tobacco leaves (Furt et al., 2010). So far, however, it has remained unclear whether and how PtdIns(4,5)P_2_ nanodomains might be formed in living cells, and what their biological functions might be.

Here, we hypothesize that spatial separation of PtdIns(4,5)P_2_ in membrane nanodomains might be a reason for the functional divergence of PtdIns(4,5)P_2_ formed by different PI4P 5-kinase isoforms in pollen tubes. We show by in vivo SD analysis that within the subapical plasma membrane of tobacco pollen tubes PtdIns(4,5)P_2_ occurs both in membrane nanodomains and in a diffuse distribution, suggesting spatially separated coexisting pools of the lipid. We further show that the functionally divergent PI4P 5-kinase AtPIP5K2 associates with plasma membrane nanodomains, whereas canonical NtPIP5K6 does not, and that nanodomain association of AtPIP5K2 is required for its divergent function as a regulator of ROP signaling and the dynamic actin cytoskeleton.

## Results

This study is aimed at understanding alternative effects of the regulatory phospholipid, PtdIns(4,5)P_2_, on processes relevant to polar tip growth, such as membrane trafficking and actin dynamics. To illustrate previously postulated alternative functions of PI4P 5-kinase isoforms within the subapical plasma membrane of pollen tubes, we performed a side-by-side comparison of phenotypic effects of overexpressing the functionally divergent pollen expressed PI4P 5-kinases, AtPIP5K2 and NtPIP5K6.

### The PI4P 5-kinases AtPIP5K2 and NtPIP5K6 exert different regulatory effects in tobacco pollen tubes

AtPIP5K2 and NtPIP5K6 are members of PI4P 5-kinase subfamily B and share a common domain architecture (Fig. 1A). Besides the parental enzymes AtPIP5K2 and NtPIP5K6 our functional comparison also included chimeric enzyme variants representing reciprocal swaps of Lin-domains termed 2swap6 and 6swap2 shown in Fig. 1A, as indicated. In growing tobacco pollen tubes, fluorescence-tagged fusions of AtPIP5K2 and NtPIP5K6 localize to the subapical plasma membrane (Fig. 1B), as previously reported for the parental enzymes and also for the chimeric variants (Stenzel et al., 2012). Please note that morphologically unaltered tubes as shown in Fig. 1B were only obtained upon very low expression levels, as in previous experiments (Stenzel et al., 2012). Despite the highly similar apparent localization in the subapical plasma membrane of the same cell (Fig. 1B), the overexpression of AtPIP5K2-EYFP or NtPIP5K6-EYFP resulted in distinct effects on cell morphology, as shown in Fig. 1C-G for the parental enzymes and for the chimeric enzyme variants 2swap6-EYFP and 6swap2-EYFP. Representative images of the respective pollen tube morphologies observed for each PI4P 5-kinase variant are shown in Fig. 1C. While untransformed control pollen tubes or pollen tubes expressing an EYFP control displayed regular pollen tube growth, overexpression of AtPIP5K2-EYFP resulted in pronounced pollen tube tip swelling (Fig. 1C). By contrast, the overexpression of NtPIP5K6-EYFP resulted in pollen tube tip branching and in extensive apical pectin deposition, here stained by ruthenium red (Fig. 1C). Please note that all pollen tubes shown in Fig. 1C and analyzed for Fig. 1D have been equally stained with ruthenium red. The overexpression of the chimeric protein 2swap6-EYFP resulted in increased apical pectin deposition (Fig. 1C), and that of the chimeric 6swap2-EYFP resulted in pollen tube tip swelling in addition to increased apical pectin deposition (Fig. 1C), indicating shifted physiological effects of the chimeric proteins. Quantifications of the degrees of pollen tube tip swelling (Fig. 1D) and of apical pectin deposition (Fig. 1E) for all expressed constructs indicate a significant difference in the morphological effects resulting from overexpressed AtPIP5K2-EYFP and NtPIP5K6-EYFP. The data furthermore indicate that the chimeric enzyme variants, in which the Lin-domains were reciprocally swapped, displayed a largely reversed pattern of morphological effects, while retaining some of the original functionality of the respective parental enzymes: When the incidence of pollen tube cell morphologies was scored in transformed populations of pollen tubes upon overexpression of the parental vs. the respective chimeric enzyme variants, overexpression of AtPIP5K2-EYFP resulted in pollen tube tip swelling and no detectable tip branching or stunted growth (Fig. 1F). By contrast, overexpression of the chimeric 2swap6-EYFP resulted in the appearance of the stunted/branched morphology concomitant with a reduced incidence of swollen cells in the population of transgenic pollen tubes (Fig. 1F). Reciprocally, overexpression of NtPIP5K6-EYFP resulted in an enhanced incidence of pollen tube tip branching and no detectable tip swelling, whereas overexpression of the chimeric 6swap2-EYFP resulted in the appearance of the swollen tip morphology concomitant with a reduced incidence of branched cells (Fig. 1G). Differences in phenotypic incidence were not due to different overexpression levels of the expressed proteins, since fluorescence intensities of transformed pollen tubes displayed no significant differences (Fig 1F and G, right panels). Please note that the effect of the RedStar-PLC-PH biosensor on pollen tube growth rates was indistinguishable from that of an expressed EYF control, and that the expression of either transgene had only a very mild effect compared to non-transformed pollen tubes (Supplemental Fig. S2).

Together, the side-by-side comparison of morphological effects of overexpressed AtPIP5K2-EYFP or NtPIP5K6-EYFP in tobacco pollen tubes suggests that these enzymes may generate PtdIns(4,5)P_2_ pools with respectively distinct regulatory effects on pollen tube growth. The determinants of such functional specification of PtdIns(4,5)P_2_ effects are not understood. The comparison of AtPIP5K2 and NtPIP5K6 effects provides the opportunity to elucidate the molecular mechanisms underlying alternative PtdIns(4,5)P_2_ functions. The cell morphologies observed upon overexpression of the chimeric enzyme variants 2swap6-EYFP and 6swap2-EYFP are consistent with a contribution of the Lin-domains in specifying the regulatory effects of PtdIns(4,5)P_2_ produced by the respective PI4P 5-kinases tested.

### PtdIns(4,5)P_2_ forms membrane nanodomains in the subapical plasma membrane of tobacco pollen tubes

The presence of a plasma membrane region enriched in PtdIns(4,5)P_2_ has previously been shown by confocal microscopy (Kost et al., 1999). To determine whether the postulated alternative effects of PtdIns(4,5)P_2_ were accompanied by further spatial segregation of PtdIns(4,5)P_2_ at a smaller scale, we analyzed the distribution of RedStar-PLC-PH, a fluorescent biosensor for PtdIns(4,5)P_2_, with high spatio-temporal resolution. In previous studies of membrane-associated protein complexes, the use of SD has enabled the observation of their lateral membrane distribution patterns at a scale resolving membrane nanodomains (Konopka et al., 2008; Sampathkumar et al., 2011; Gadeyne et al., 2014; Bloch et al., 2016; Synek et al., 2017). Therefore, we used SD to analyze the distribution of RedStar-PLC-PH in pollen tubes (Fig. 2). In a median confocal section of a pollen tube expressing RedStar-PLC-PH fluorescence was present only at the plasma membrane and possibly at the immediate periphery, but not in the volume of the cell, as indicated by confocal z-stacks projected in xy, yz and xz orientations (Fig. 2A, recording planes indicated by arrowheads). A progression through the z-stack also illustrates that the volume of the pollen tube was devoid of RedStar-PLC-PH fluorescence (Fig. 2B, Supplemental Movie 1), whereas a spotty fluorescence distribution was observed at the cell surface. A 3D-projection of the same image-stack clearly shows the presence of numerous punctate signals in addition to diffuse fluorescence in the plane of the membrane at the cell surface (Fig. 2C, Supplemental Movie 2), indicating possible subcompartmentalization of PtdIns(4,5)P_2_ in the plasma membrane of pollen tubes. The plasma membrane nanodomains decorated by RedStar-PLC-PH (from now also referred to as “PtdIns(4,5)P_2_ nanodomains”) occurred in a region between 1-12 µm distant from the tips of both growing and non-growing pollen tubes, with 50% of the nanodomains concentrating between 3 and 6 µm distance from the tip (Fig. 2D). PtdIns(4,5)P_2_ nanodomains of different sizes were found uniformly distributed with increasing distance from the pollen tube tips and with no difference between growing or non-growing tubes (Fig. 2E). The mean apparent diameters of the PtdIns(4,5)P_2_ nanodomains were 0.7 ± 0.1 µm in growing and 0.6 ± 0.1 µm in non-growing pollen tubes (Fig. 2F), with the majority of nanodomains displaying diameters of approx. 0.6 µm in both growing and non-growing pollen tubes (Fig. 2G). Detailed scrutiny of the RedStar-PLC-PH signal intensities quantified along the dashed lines in Fig. 2H and Fig. 2I indicated that the intensity of nanodomains decorated by the PtdIns(4,5)P_2_ reporter was roughly two-fold higher than the diffuse reporter fluorescence in the membrane (Fig. 2H, I), which by itself was substantially above background (indicated by dashed lines in the intensity profiles in Figs. 2H and I). The SD analysis of pollen tubes expressing the RedStar-PLC-PH biosensor for PtdIns(4,5)P_2_ thus indicates the presence of PtdIns(4,5)P_2_ nanodomains in addition to diffuse reporter fluorescence within the subapical plasma membrane in a pattern consistent with spatially separated coexisting pools of PtdIns(4,5)P_2_.

**Figure 2.**
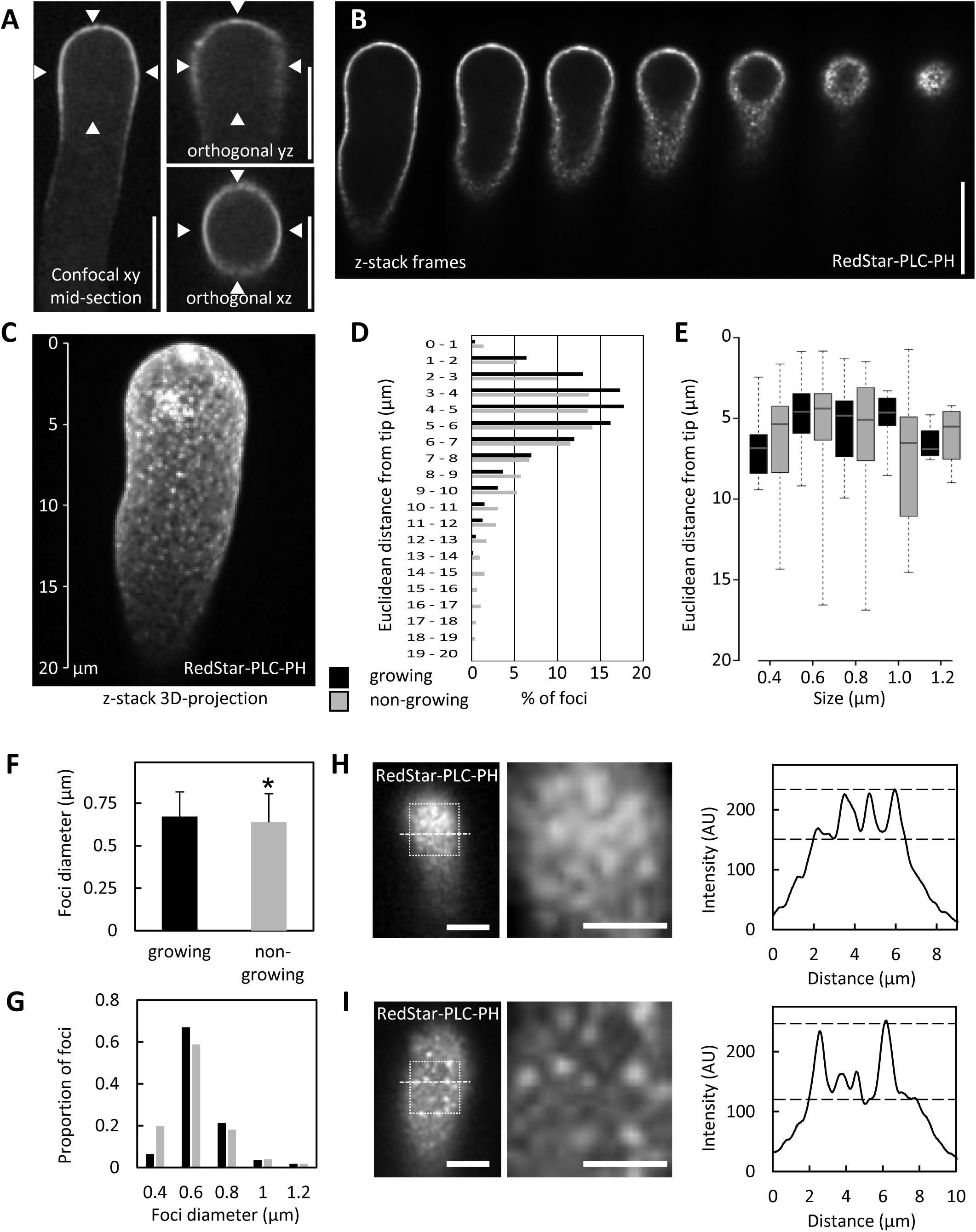
A PtdIns(4,5)P_2_ biosensor decorates membrane nanodomains in the subapical plasma membrane of tobacco pollen tubes. The fluorescence distribution of the PtdIns(4,5)P_2_-specific reporter RedStar-PLC-PH was monitored upon transient expression in tobacco pollen tubes by SD. **A**, Projections of confocal z-stacks with xy, yz and xz orientations of a representative pollen tube expressing RedStar-PLC-PH. Scale bars, 10 µm. **B**, Sequence of images extracted from SD *z*-stack acquisition of tobacco pollen tubes expressing RedStar-PLC-PH (from median confocal plane to the cell surface). PtdIns(4,5)P_2_ nanodomains are visible as bright dots at the cell surface. Please note the absence of fluorescence at the cell interior. Scale bar, 10 µm. **C**, 3D projection from a *z*-stack acquisition of a representative pollen tube expressing RedStar-PLC-PH. Scale as indicated. **D**, Distribution of PtdIns(4,5)P_2_ nanodomain over the euclidean distance from the pollen tube tip, as determined for growing pollen tubes (black bars) or for non.-growing pollen tubes grey bars). The analysis is based on 8881 nanodomains from 8 cells (growing) or 17204 nanomonains from 10 cells (non-growing). **E**, Size distribution of nanodomains decorated by RedStar-PLC-PH along the tips of growing (black bars) or non-growing pollen tubes (grey bars) over the Euclidean distance from the pollen tube tips. Data represent 221 nanodomains from 8 cells (growing); or 227 nanodomains from 10 cells (non-growing), respectively. **F**, Mean nanodomain diameters were calculated for growing (black bar) or non-growing pollen tubes (grey bar) form the data shown in (E). Data are given as means ± standard deviation. The asterisk indicates a significant difference according to a Student’s T-test (*, p≤0.05). **G**, The distribution of size categories of nanodomain diameters was scored in growing (black) and in non-growing pollen tubes (grey). The distribution was calculated from the same data as shown in (F). **H, I**, The intensities of PtdIns(4,5)P_2_ nanodomains decorated by RedStar-PLC-PH were recorded by SD in cell surface scans of growing (H) or non-growing pollen tubes (I). Left panels, Representative SD images of the cell surface. Scale bars, 5 µm. White boxes, regions of interest enlarged in middle panels (scale bars there, 3 µm). Right panels, Fluorescence intensity plots derived from dashed lines in left panels. Dashed lines in the intensity plots, estimated levels of diffuse marker fluorescence (lower line) and of nanodomain fluorescence (upper line).

### PtdIns(4,5)P_2_ nanodomains are highly dynamic

Punctate signals of the the RedStar-PLC-PH biosensor were monitored over time at the cell surface of growing and non-growing pollen tubes (Fig. 3A, B, Supplemental Movies 3-6). A kymograph analysis (Fig. 3C) indicates dynamic behavior of the nanodomains with plasma membrane lifetimes between few seconds to few minutes (Fig. 3D). In growing pollen tubes, approx. 75% of the PtdIns(4,5)P_2_ nanodomains exhibited lifetimes below 10 s (Fig. 3D). While approx. 65% of PtdIns(4,5)P_2_ nanodomains in non-growing pollen tubes also had lifetimes below 10 s, the non-growing cells additionally displayed a population of PtdIns(4,5)P_2_ nanodomains with longer lifetimes of up to 2 min (Fig. 3D). The incidence of PtdIns(4,5)P_2_ nanodomain lifetimes above 60 s was found to be approx. 4-times higher in non-growing than in growing cells (Fig. 3D inset). An analysis of the lifetimes of PtdIns(4,5)P_2_ nanodomains at different distances from the pollen tube tip indicates that nanodomains with longer lifetimes occurred mostly further away from the tip in non-growing pollen tubes, while the majority of the nanodomains with shorter lifetimes occurred significantly closer to the tip in growing pollen tubes (Fig. 3E). The dynamic pattern of plasma membrane nanodomains decorated by the RedStar-PLC-PH biosensor is consistent with perpetual competitive recruitment of proteins to the nanodomains - possibly displacing the biosensor - and/or represents an active balance of biosynthesis by PI4P 5-kinases and breakdown of PtdIns(4,5)P_2_.

**Figure 3.**
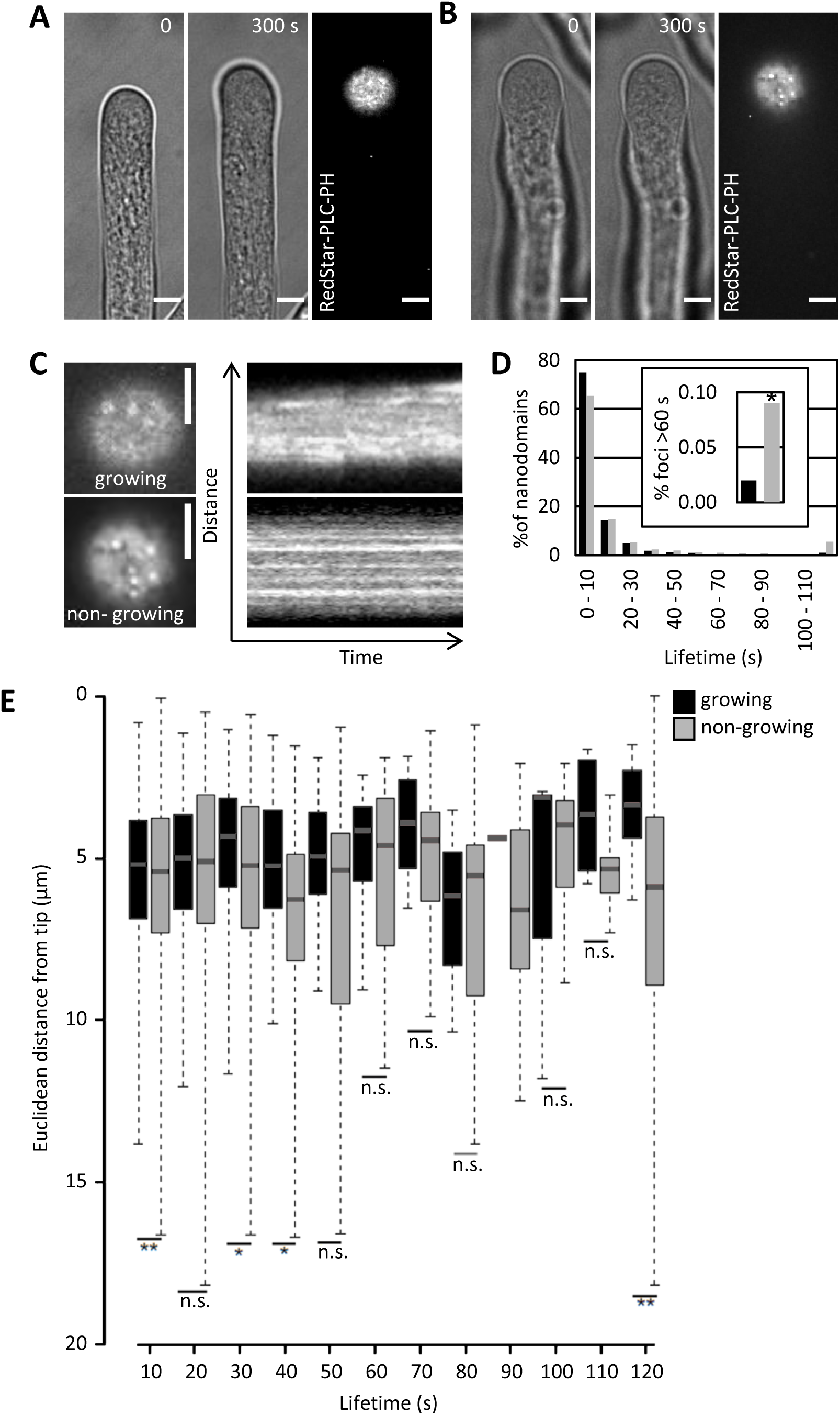
RedStar-PLC-PH-labeled PtdIns(4,5)P_2_ nanodomains are dynamic. The dynamics of PtdIns(4,5)P_2_ nanodomains at the cell surface were recorded by SD in tobacco pollen tubes upon transient expression of the RedStar-PLC-PH biosensor from the Lat52 promoter. **A, B**, PtdIns(4,5)P_2_ nanodomains in a growing (A) or non-growing tobacco pollen tubes (B). Left panels, Growth or no growth of the pollen tube according to bright field image frames from image acquisition over a period of 2 min at 0.5 frames s^-1^; only the first and last frames are shown. Right panels, SD imaging of the last frames from the same pollen tubes with the confocal plane focused at the PtdIns(4,5)P_2_ nanodomains at the cell surface. Scale bars, 5 µm. **C**, Kymograph analysis of the image sequences obtained for growing (upper panels) or non-growing cells (lower panels). Left, fluorescence images at the cell surface (magnified from images in panels A and B; Scale bars, 5 µm. Right panels, Kymographs from the same pollen tubes over a period of 5 min. **D**, Life time distribution of dynamic PtdIns(4,5)P_2_ nanodomains decorated by RedStar-PLC-PH in growing (black bars) or non-growing pollen tubes (grey bars). Data represent 1415 tracks from eight cells (growing) or 3203 tracks from 16 cells (non-growing pollen tubes). Inset, Total percentage of nanodomains with lifetimes between 60 and 120 s in growing (black bar) or non-growing pollen tubes (grey bar). **E**, Distribution of lifetimes of PtdIns(4,5)P_2_ nanodomains over the Euclidean distance from the tips of growing (black bars) or non-growing pollen tubes (grey bars). Asterisks indicates significant differences between the corresponding values (lifetimes or distances) according to a Student’s T-test (**, p≤0.01; *, p≤0.05; n.s., not significant).

### AtPIP5K2 occupies nanodomains whereas NtPIP5K6 displays uniform distribution in the plasma membrane of pollen tubes

To assess the biosynthesis of PtdIns(4,5)P_2_ in this context, we monitored the lateral distribution of fluorescence-tagged variants of AtPIP5K2-EYFP, NtPIP5K6-EYFP or the chimeric variants 2swap6-EYFP and 6swap2-EYFP (see Fig. 1A) in pollen tubes by SD (Fig. 4). All constructs were expressed from the Lat52 promoter. Median confocal sections indicated association with the plasma membrane in all cases (Fig 4A-D, left panels). When confocal z-stacks were recorded and the frames representing the plasma membrane surface were visually examined, AtPIP5K2-EYFP associated with plasma membrane nanodomains (Fig. 4A, middle and right images, Supplemental Movies 3 and 4), whereas NtPIP5K6-EYFP displayed a more uniform distribution (Fig. 4B, middle and right images). The substantial difference in the association of AtPIP5K2-EYFP with nanodomains vs. the uniform distribution of NtPIP5K6-EYFP was evident from the intensity plots (far right) recorded along the dashed lines in the detail views of the surface scans, as indicated. When the plasma membrane distribution of AtPIP5K2-EYFP was compared side-by-side to that AtPIP5K5-EYFP, which causes apical pectin secretion and pollen tube branching upon overexpression in pollen tubes (Ischebeck et al., 2008; Ischebeck et al., 2010b), a similar difference in distribution patterns was observed by SD, with AtPIP5K2-EYFP in nanodomains and AtPIP5K5-EYFP displaying diffuse distribution similar to that observed for NtPIP5K6-EYFP (Supplemental Fig. S3). The data indicate that the pollen tube tip swelling morphology is associated with AtPIP5K2-EYFP located in plasma membrane nanodomains, whereas “secretion phenotypes” are associated with NtPIP5K6-EYFP or AtPIP5K5-EYFP which display a continuous plasma membrane distribution.

**Figure 4.**
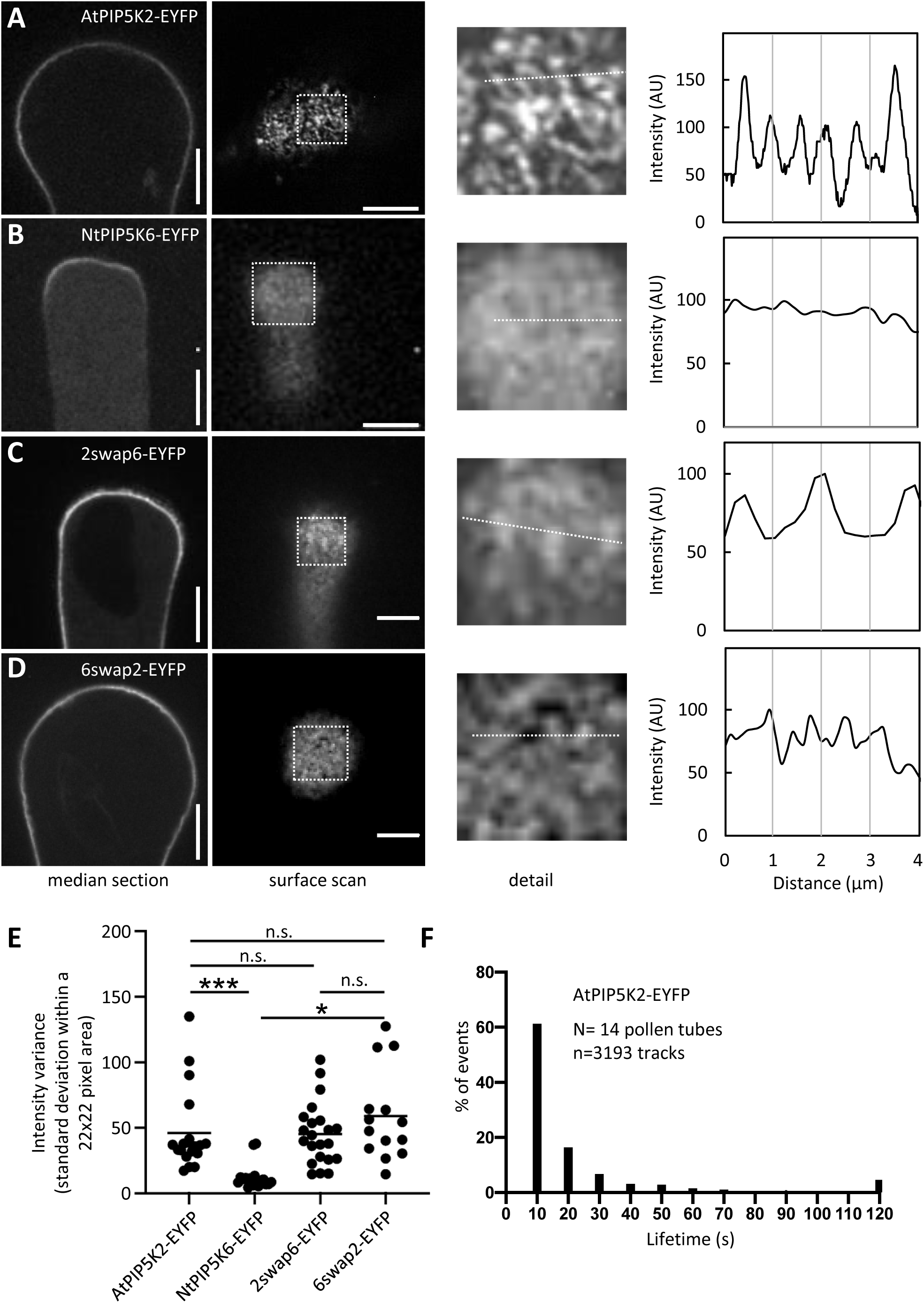
AtPIP5K2-EYFP occupies nanodomains whereas NtPIP5K6-EYFP displays uniform distribution in the plasma membrane. The fluorescence distribution of AtPIP5K2-YFP, NtPIP5K6-YFP, or their chimeric variants 2swap6-EYFP or 6swap2-EYFP at the plasma membrane was monitored by SD upon transient expression in tobacco pollen tubes from the Lat52 promoter. **A**, AtPIP5K2-EYFP; **B**, NtPIP5K6-EYFP; **C**, 2swap6-EYFP; **D**, 6swap2-EYFP. Left images, Median confocal sections of representative pollen tubes showing the fluorescence signal at the plasma membrane. Center images, Surface scan of the same cells extracted from z-stack acquisition. White boxes, Areas of interest enlarged in right images, giving a detail view of the distribution of the fusion proteins at the plasma membrane. Far right, Fluorescence intensity plots derived from the dashed lines in the magnified detail images. Scale bars, 5 µm. **E**, Non-biased assessment of uniformity of fluorescence intensity distributions according to the standard deviations of overall fluorescence intensity values. The standard deviations of all intensity values in a 22×22 pixel area of an image were calculated as a measure of uniformity for cells expressing AtPIP5K2-EYFP (N=19), NtPIP5K6-EYFP (N=17), 2swap6-EYFP (N=22) and 6swap2-EYFP (N=14). Asterisks indicate significant differences according to a Student’s T-test (***, p≤0.001; *, p≤0.05; n.s., not significant). **F**, Lifetime distribution of AtPIP5K2-EYFP nanodomains. Pollen tubes expressing AtPIP5K2-YFP were imaged by SD at a frame rate of 0.5 frames s^-1^ for 2 min. The distribution of lifetimes is given based on 31193 tracks recorded from 14 cells.

To assess the role of the Lin-domains in mediating nanodomain vs. diffuse membrane patterns, we analyzed the distribution of the chimeric proteins 2swap6-EYFP and 6swap2-EYFP. The 2swap6-EYFP chimera displayed a clear reduction in the degree of nanodomain association compared to the parental AtPIP5K2-EYFP protein, as indicated by the surface scan and the corresponding line intensity plot (Fig. 4C). The exchange of the Lin domain of AtPIP5K2 for that of NtPIP5K6 did not fully abolish nanodomain association of the 2swap6-EYFP fusion, however, possibly because of additional features of the AtPIP5K2 protein which may still favour nanodomain association. The reciprocal 6swap2-EYFP chimera displayed substantially enhanced association with membrane nanodomains when compared to the uniform distribution of the parental NtPIP5K6-EYFP protein, as indicated by the surface scan and line intensity plot (Fig. 4D). The distribution of the 6swap2-EYFP fusion shows that the Lin-domain of AtPIP5K2 can confer a significant degree of nanodomain association to the NtPIP5K6-EYFP protein.

As the comparison of line intensity patterns might be subject to experimental bias (e.g., through the manual positioning of the dashed line), we applied a non-biased analysis based on the standard deviations of pixels intensity values from multiple full images recorded by SD at the cell surface. High standard deviations among intensity values reflect the existence of a heterogenous distribution of fluorescence intensity values, and therefore of nanodomains at the membrane. By contrast, low standard deviations reflect a homogeneous distribution of intensity values and therefore of a diffuse distribution of the fluorescence at the membrane. The standard deviations for surface scans of pollen tubes expressing AtPIP5K2-EYFP, NtPIP5K6-EYFP, 2swap6-EYFP or 6swap2-EYFP are given in Fig. 4E and correspond well with the data from the visual examination and from the line intensity plots.

The plasma membrane nanodomains decorated by AtPIP5K2-EYFP were dynamic (Supplemental Movies 7-8) and the analysis of their lifetime distribution (Fig. 4F) resembled that of RedStar-PLC-PH nanodomains (see Fig 2D), with more than 60% of nanodomains having lifetimes shorter than 10 s. Interestingly, we observed a small percentage (approx. 5 %) of AtPIP5K2-EYFP nanodomains with longer lifetimes (>2 min) in a similar pattern as with RedStar-PLC-PH nanodomains from non-growing pollen tubes (see Fig 2D).

Overall, the different lateral plasma membrane distributions of AtPIP5K2-EYFP and NtPIP5K6-EYFP appear to correlate with the divergent respective functionalities of the enzymes. The divergent membrane distribution patterns support the notion that subcompartmentalization of PtdIns(4,5)P_2_ production by these enzymes may underlie the functional specification of PtdIns(4,5)P_2_ regulatory effects. The data furthermore support a role for the Lin-domain as one factor contributing to nanodomain association or uniform membrane association of PI4P 5-kinases, thereby directing PI4P 5-kinases into different regulatory contexts.

### AtPIP5K2-EYFP and PtdIns(4,5)P_2_ nanodomains occur in a spatially correlated pattern at the plasma membrane

The results so far indicated the presence of plasma membrane nanodomains with similar dynamics for PtdIns(4,5)P_2_ and AtPIP5K2-EYFP. Therefore, we next characterized the spatial distribution of nanodomains occupied by AtPIP5K2-EYFP at the plasma membrane in more detail. Confocal z-stacks acquired by SD indicate that AtPIP5K2-EYFP fluorescence was limited to the cell surface, with little or no fluorescence in the volume of the cell (Fig. 5A, left panel). A 3D-projection of the z-stack shows the presence of punctate signals at the cell surface (Fig. 5A, right panel). Membrane-association of the AtPIP5K2-EYFP marker was assessed by costaining with the lipophilic dye, FM 4-64 (Fig. 5 B, C). While the distribution of AtPIP5K2-EYFP coincided with that of FM 4-64 (Fig. 5C), we noted a small but reproducible right shift of the AtPIP5K2-EYFP signal (Fig. 5C), suggesting that AtPIP5K2-EYFP occupied a peripheral region further inside the cell than the plasma membrane dye. This result is further supported by the patterns observed upon coexpression of AtPIP5K2-EYFP together with the PtdIns(4,5)P_2_ biosensor RedStar-PLC-PH, where we also observed a small spatial distinction with the AtPIP5K2-EYFP marker localizing further inward from the plasma membrane into the cell periphery (Fig. 5D).

**Figure 5.**
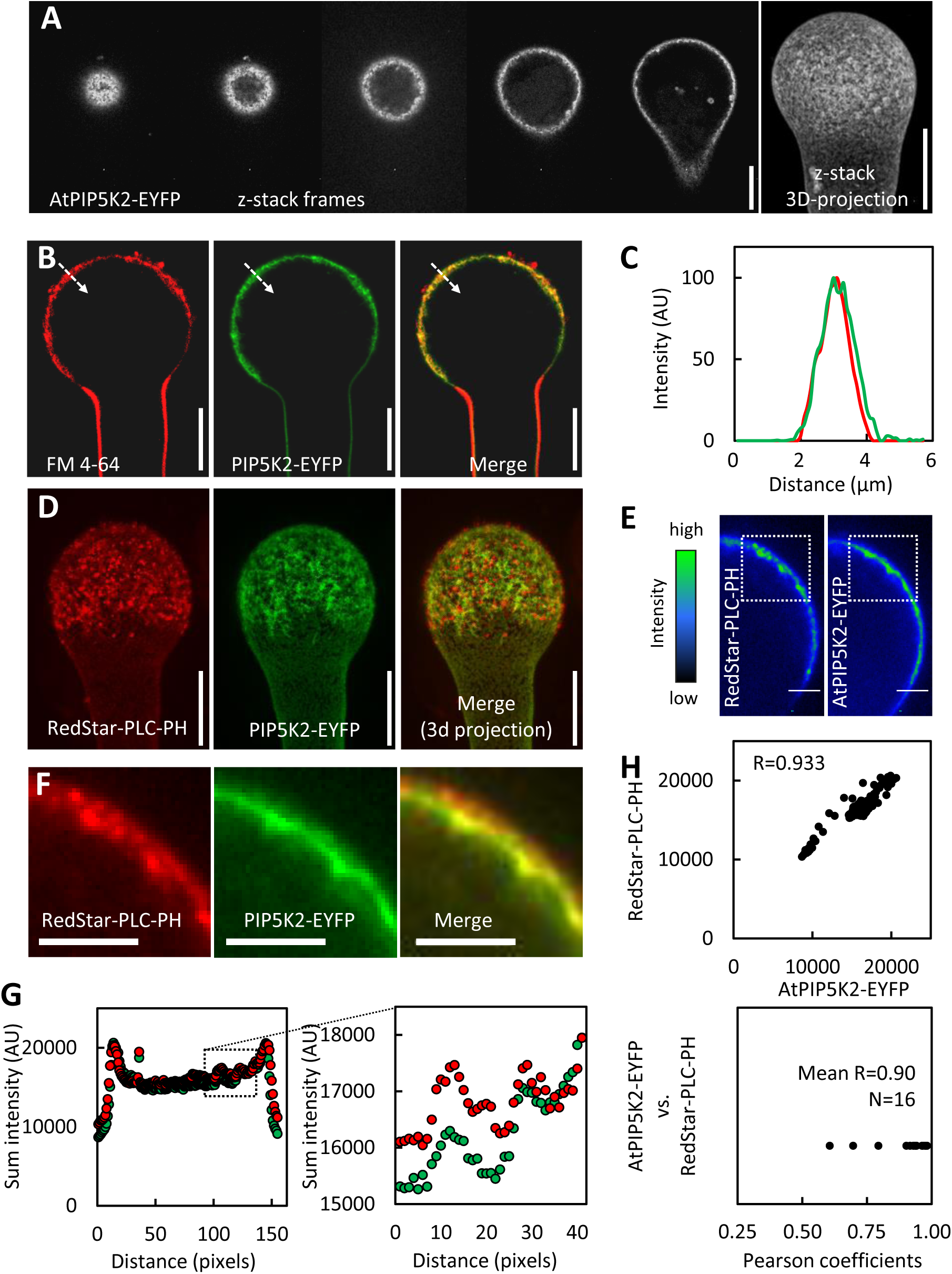
AtPIP5K2-EYFP and PtdIns(4,5)P_2_ nanodomains occur in a spatially correlated pattern at the plasma membrane. The fluorescence distribution of AtPIP5K2-YFP and RedStar-PLC-PH at the plasma mebrane were observed by SD or laser-scanning confocal microscopy (LSM) upon transient expression in tobacco pollen tubes from the Lat52 promoter. **A**, Representative sequence of images extracted from SD z-stack acquisition of a tobacco pollen tube expressing AtPIP5K2-YFP (left to right, from cell surface to median confocal section). Right image, 3D projection obtained from the same full z-stack acquisition. AtPIP5K2-YFP nanodomains are visible as bright dots at the cell surface. Scale bars, 5 µm. **B**, Representative LSM images of a tobacco pollen tube expressing AtPIP5K2-YFP and costained with the membrane dye, FM 4-64, as indicated. Scale bars, 10 µm. **C**, Fluorescence intensity plot derived from the dashed line across the plasma membrane, as indicated in (B). **D**, A representative 3D reconstruction from SD z-stack acquisition of a pollen tube coexpressing the PtdIns(4,5)P_2_ biosensor RedStar-PLC-PH and AtPIP5K2, as indicated. Right panel, merged image. Scale bar, 10 µm. **E**, Representative median confocal frames extracted from the z-stack projection shown in (D) and representing the individual channels for RedStar-PLC-PH and AtPIP5K2-EYFP with heatmap colors, as indicated. Note that high intensity values from both channels localize in the same areas at the cell periphery. **F**, Detail fluorescence micrographs of the regions of interest indicated by the white boxes in panel (E), as indicated, which were unsed for the sum intesity analysis. **G**, The sum intensities were determined for the vertical pixel columns of each channel from images in panel F (please refer to Supplementary Fig. S4 and Materials and Methods for details). Left panel, Sum intensities for RedStar-PLC-PH (red) and AtPIP5K2-YFP (green); right panel, Detail view if the region of interest indicated by the box in the left panel. **H**, Correlation analysis of sum intensity profiles obtained for RedStar-PLC-PH and AtPIP5K2-EYFP. Upper panel, Representative correlation plot for the fluorescence distributions given in panel (G). The correlation is characterized by a high Pearson coefficient R of 0.933. Lower panel, Pearson coefficients obtained accordingly from 16 correlation analyses, with a mean R of 0.9, indicating consistently high correlations between the söatial distributions of RedStar-PLC-PH and AtPIP5K2-EYFP.

Both the PtdIns(4,5)P_2_-biosensor and the AtPIP5K2-EYFP marker displayed membrane nanodomain association at the cell surface when analyzed by SD, so we next asked whether the distribution patterns of these markers were spatially correlated. To assess the relative distribution patterns of AtPIP5K2-EYFP and RedStar-PLC-PH, we attempted to analyze flattened images from 3D-projections of cells coexpressing both markers, such as shown in Fig. 5D. Due to the density of the signals, however, colocalizing signals in these analyses could not be distinguished from random coincidence. To nonetheless provide a measure of the relative positioning of AtPIP5K2-EYFP and RedStar-PLC-PH distributions, we resorted to the quantitative analysis of median confocal sections (Fig. 5E), which display lesser complexity as shown in Fig. 5 E-I. Fluorescence signals from multiple stacks were analyzed by adding the intensity values of the vertical pixel columns for each fluorescence channel (AtPIP5K2-EYFP and RedStar-PLC-PH in Fig. 5E; for details, please see the Material and Methods section and Supplemental Fig. S4). Detail patterns for the individual fluorescence channels are given for a representative pollen tube in Fig. 5F. The resulting sum intensity profiles (Fig. 5G) provide a measure for the intensity distribution along the curved plasma membrane surface of the pollen tube cells. The sum intensity profiles thus obtained for AtPIP5K2-EYFP and for RedStar-PLC-PH display similar detail patterns (Fig. 5G, right panel). Correlation analysis of the fluorescence patterns, which can be numerically assessed by calculating the corresponding Pearson coefficient R, which can range from 0 (no correlation) to 1 (perfect correlation), was performed and is shown for one representative pollen tube in Fig. 5H (upper panel; R=0.933). The analysis of pollen tube cells coexpressing both markers (16 cells, each with five individual measurements) consistently indicates a tight correlation of the markers with a mean R-value of 0.90 (Fig. 5H, lower panel).

Together, the data indicate that AtPIP5K2-EYFP distributes in the plasma membrane in a nanodomain pattern that is spatially correlated to the positioning of the PtdIns(4,5)P_2_-biosensor and additionally reaches inward towards the cytoplasmic periphery.

### AtPIP5K2-EYFP can associate with filamentous structures underlying the plasma membrane

As pollen tube tip swelling morphology, such as that observed upon overexpression of AtPIP5K2-EYFP (Fig. 1C-G), has previously been linked to changes in the actin cytoskeleton (Klahre et al., 2006; Klahre and Kost, 2006; Kost, 2008; Ischebeck et al., 2011), we further investigated the distribution of AtPIP5K2-EYFP in relation to pollen tube actin (Fig. 6). A link to cytoskeletal control is supported by the observation that in some cells overexpressing AtPIP5K2-EYFP we observed not only an inward-shift in the localization but to varying degrees also the formation of a mesh-like pattern underlying the plasma membrane (Fig. 6A). While this pattern was only observed in cells overexpressing AtPIP5K2 and likely does not represent the physiological localization of AtPIP5K2, it might nonetheless indicate a principal capability of the enzyme to associate with membranes or other structures underlying the plasma membrane. The mesh-like pattern is inward oriented when compared to the distribution of PtdIns(4,5)P_2_ biosensor RedStar-PLC-PH (Fig. 6A), as can be seen from the detail micrographs in Fig. 6A, and suggests a cortical cytoskeletal structure. To determine the distribution of AtPIP5K2-EYFP relative to actin, we next analyzed AtPIP5K2-EYFP together with the coexpressed actin marker, LifeAct-mCherry (Fig. 6B). While no colocalization of the markers was found, a coinciding pattern was observed at the interface of actin filaments with the plasma membrane, where nanodomain-associated AtPIP5K2-EYFP displayed increased intensity (Fig. 6B, upper panels) or even protruded inwards towards the filaments (Fig. 6B, lower panels, arrowheads in the detail panel). Please note that in these experiments both the distribution of AtPIP5K2-EYFP and that of the LifeAct-mCherry may be influenced by the overexpression of the AtPIP5K2-EYFP marker, which results in tip swelling, and that in normal-growing pollen tubes, actin will not protrude into the tip region of the pollen tubes. A pattern of relative distributions similar to that seen in Fig. 6B was also observed when the chimeric 6swap2-EYFP marker was coexpressed with LifeAct-mCherry (Fig. 6C), with 6swap2-EYFP fluorescence protruding inward from the plasma membrane (Fig. 6C, arrowheads in the detail panel). These data suggest that the Lin-domain of AtPIP5K2 is sufficient to mediate the association of the chimeric enzyme with plasma membrane regions involved in the peripheral membrane attachment of actin filaments. The inward-oriented protrusions decorated by AtPIP5K2-EYFP or by 6swap2-EYFP (Fig. 6B, C) might represent plasma membrane pulled towards the inside of the cell, possibly through enhanced membrane attachment of actin filaments. A truncated variant of AtPIP5K2-EYFP in which the N-terminal NT-, MORN- and Lin-domains were deleted (AtPIP5K2_344-754_-EYFP) lost association with plasma membrane nanodomains and decorated a mesh-like pattern at the cell periphery, which possibly reflects association with cortical actin (Fig. 6D). This observation suggests that features of the AtPIP5K2 protein other than the deleted domains are responsible for an association of the enzyme with the actin cytoskeleton, and that in the absence of the membrane-associating Lin-domain this unknown element may mediate association of the truncated protein with actin structures within the cell.

**Figure 6.**
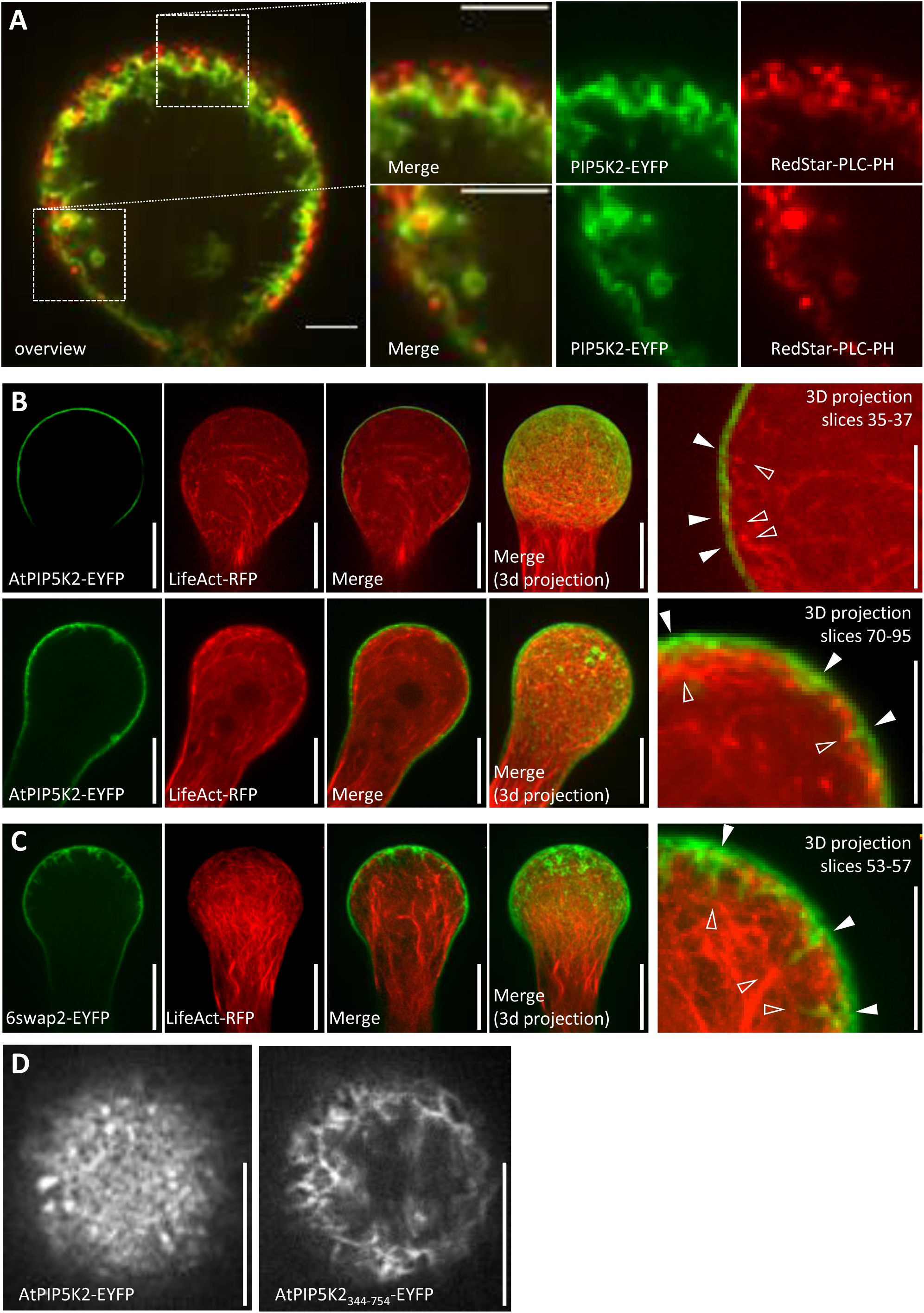
AtPIP5K2-EYFP can associate with filamentous structures underlying the plasma membrane. The fluorescence distribution of AtPIP5K2-YFP, RedStar-PLC-PH and Lifeact-RFP was monitored by SD upon transient expression in tobacco pollen tubes from the Lat52 promoter. In approximately 25 % of cells with high fluorescence intensities, the AtPIP5K2-EYFP marker decorated a mesh-like structure underlying the plane of signals decorated by RedStar-PLC-PH nanodomains. **A**, Median cross section of the swollen tip of a pollen tube coexpressing AtPIP5K2-EYFP (green) and RedStar-PLC-PH (Red). Only the merged overview image is shown. White boxes, Regions of interest detailed in the right panels showing individual fluorescence channels channels and a merged image, respectively. Scale bars, 5 µm. **B, C**, The distribution of AtPIP5K2-EYFP (B) or the chimeric 6swap2-EYFP (C) was monitored in tobacco pollen tubes relative to thath of coexpressed LifeAct-RFP, an in vivo marker for actin. Median confocal sections are shown for the individual fluorescence channels and the merged images, together with 3D reconstructions from SD z-stack acquisition, as indicated. Far right, magnified view of maximum intensity projections of several z-stack frames as indicated, highlighting AtPIP5K2-EYFP fluorescence at the plasma membrane (green) coinciding with peripheral actin structures (red), as indicated by the arrowheads (closed, AtPIPP5K2-EYFP; open, LifeAct-RFP). **B**, upper panels, A representative swollen pollen tube displaying AtPIP5K2-EYFP mostly at the cell surface. **B**, lower panels, Pollen tube displaying AtPIP5K2-EYFP in a pattern also decorating the are from the plasma membrane inward. **C**, Pollen tube displaying 6swap2-EYFP in a pattern also decorating the area from the plasma membrane inward. **D**, The distribution patterns of AtPIP5K2-EYFP and an N-terminally truncated variant AtPIP5K2_344-754_-EYFP were monitored by SD at the cell surface of tobacco pollen tubes upon transient expression from the Lat52 promoter. Please note the association of the truncated AtPIP5K2_344-754_-EYFP fusion with a mesh-like pattern underlying the plasma membrane. The pattern has consistently been observed in 12 out of 12 transformed cells. Scale bars, 10 µm.

The combined observations suggest a facultative spatial association of AtPIP5K2-EYFP with membrane regions involved in actin attachment. This notion is consistent with an effect of AtPIP5K2-EYFP on actin to mediate tip swelling, similar to previous reports (Klahre et al., 2006; Klahre and Kost, 2006; Kost, 2008; Ischebeck et al., 2011). The data furthermore support a role for the Lin-domain of AtPIP5K2 as one main factor in positioning the enzyme and PtdIns(4,5)P_2_ for a possible regulatory effect on actin. As such a regulatory effect of PtdIns(4,5)P_2_ has not previously been shown for plant cells, further experiments were performed to address its effects on actin distribution in more detail.

### AtPIP5K2-EYFP controls the dynamic actin cytoskeleton in pollen tubes

To assess regulation of the pollen tube actin cytoskeleton by AtPIP5K2-EYFP, we monitored actin in pollen tubes coexpressing variants of AtPIP5K2 or NtPIP5K6 together with the in vivo actin markers LifeAct-mCherry (Riedl et al., 2008; Vidali et al., 2009; Lichius and Read, 2010) or mTalin-YFP (Kost et al., 1998; Fu et al., 2001) (Fig. 7). The distribution of actin in the pollen tube tip of cells coexpressing PI4P 5-kinase variants with LifeAct-mCherry was visually examined in confocal z-stack obtained by SD (Fig. 7A, as indicated) and was quantitatively assessed first according to the occupancy of the actin signal in the apical region of the cells (Fig. 7B). Pollen tubes coexpressing LifeAct-RFP with an EYFP control did not have detectable actin structures in the apical region of the cells (Fig. 7A, as indicated). By contrast, cells overexpressing AtPIP5K2-EYFP and displaying tip swelling were characterized by an increased abundance of actin filaments in the apical region (Fig. 7A, as indicated). Differences were highly significant according to the actin occupancy measurements (Fig. 7B). Pollen tubes overexpressing NtPIP5K6-EYFP or the 2swap6-EYFP chimera displayed no or only weak tip swelling, respectively, (Fig. 7A, as indicated) and displayed significantly less actin occupancy in the tip compared to AtPIP5K2-EYFP (Fig. 7B). Tip swelling was observed upon overexpression of the 6swap2-EYFP chimera (Fig. 7A, as indicated) and was accompanied by a significantly increased apical actin occupancy compared to the pattern observed with the parental NtPIP5K6-EYFP (Fig. 7B). In these experiments, the membrane association of the PI4P 5-kinase variants did not differ, as determined by the ratios of plasma membrane/cytosolic fluorescence intensities in median confocal planes observed for the different markers, as indicated (Fig. 7C).

**Figure 7.**
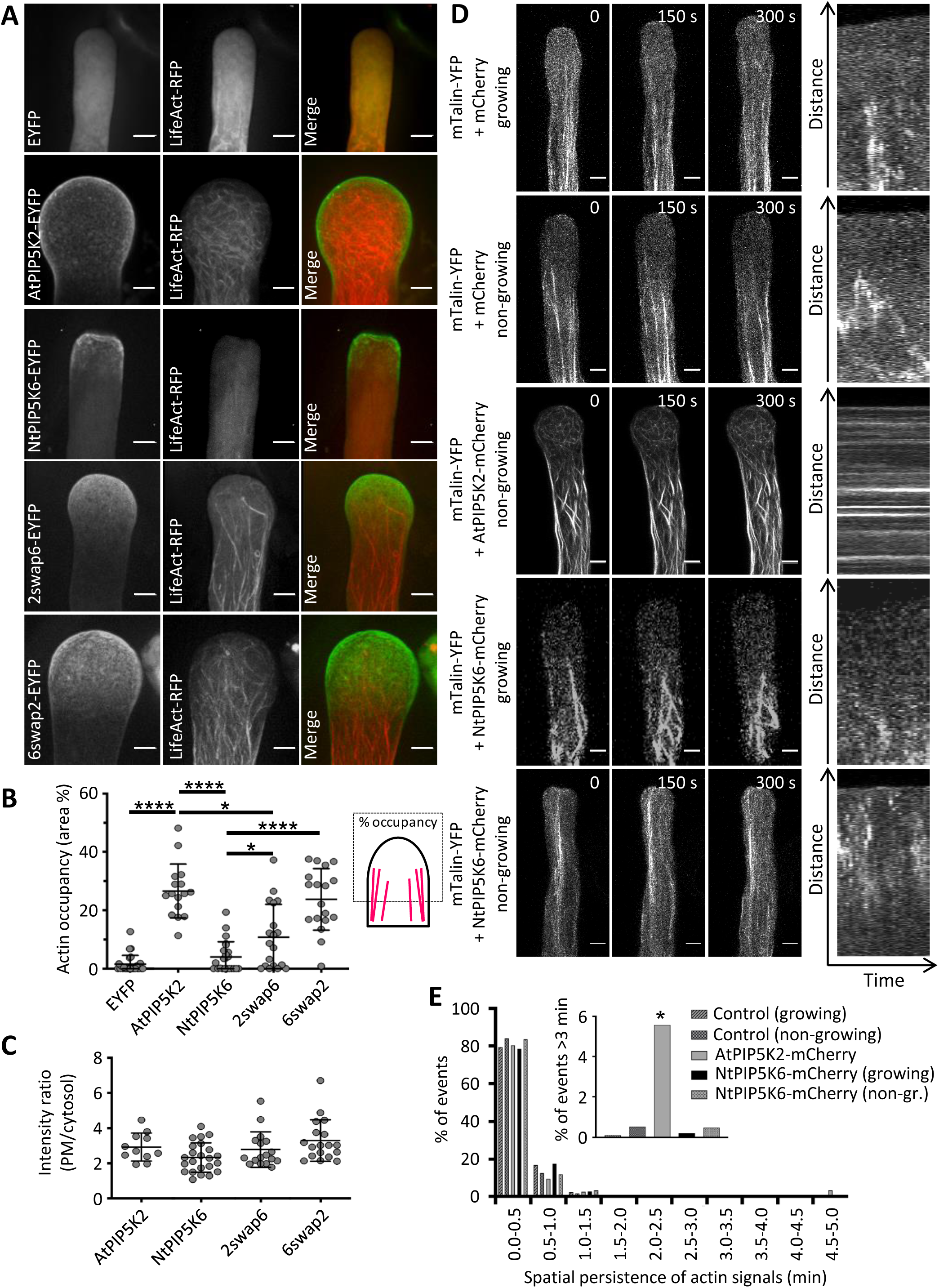
AtPIP5K2-EYFP controls the dynamic actin cytoskeleton of pollen tubes. The actin cytoskeleton was monitored in vivo by SD in tobacco pollen tubes upon coexpression of LifeAct-RFP respectively with an EYFP control, AtPIP5K2-EYFP, NtPIP5K6-EYFP, 2swap6-EFP or 6swap2-EYFP, as indicated. **A**, Representative 3D reconstructions from z-stack acquisition are shown for the individual fluorescence channels and the merge, as indicated. Scale bars, 5 µm. **B**, Occupancy of the LifeAct-RFP signal in the pollen tube tips. Data are from 26 cells (LifeAct-RFP/EYFP); 16 cells (LifeAct-RFP/AtPIP5K2-EYFP); 23 cells (LifeAct-RFP/NtPIP5K6-EYFP) 18 cells (LifeAct-RFP/2swap6-EYFP); or 19 cells (LifeAct-RFP/6swap2-EYFP), respectively. Asterisks indicate significant differences according to a Student’s T-test, as indicated (****, p≤0.0001; *, p≤0.05). **C**, Membrane association of the PI4P 5-kinase variants was assessed by calculating the ratio of plasma membrane (PM)-associated fluorescence and cytosolic fluorescence from the images used for the analysis in (B), with no statistical differences between the distributions. **D**, The dynamics of the actin cytoskeleton was monitored in tobacco pollen tubes coexpressing the in vivo actin marker mTalin-YFP with an mCherry control, AtPIP5K2-mCherry, NtPIP5K6-mCherry, 2swap6-mCherry or 6swap2-mCherry, as indicated. Time series of z-stacks were acquired by LSM at 0.1 frames s^-1^ over a period of five min. Images in the left panels show representative maximum intensity projections of the mTalin-YFP channel of pollen tubes observed at t=0, t=150s and t=300s for growing or non-growing pollen tubes, as indicated. Right panels, Representative kymograph analyses of the mTalin-YFP channel. Please note that horizontal lines indicate structures labeled by mTalin-YFP that remained static over the recorded period. Scale bars, 5 µm. **E**, The spatial persistence of the mTalin-YFP fluorescence was extracted from kymographs and plotted (for details of this analysis, please also see Materials and Methods section). Data represent 12 cells (mTalin-YFP/mCherry, growing); 10 cells (mTalin-YFP/mcherry, non-growing); 21 cells (mTalin-YFP/AtPIP5K2-mcherry); 7 cells (mTalin-YFP/ NtPIP5K6-mCherry growing pollen tubes); 15 cells (mTalin-YFP/NtPIP5K6-mCherry, not growing). Inset, The percentage of mTalin-YFP kymograph signals persisting for period >3 min were plotted, and a significantly higher incidence of persisting signals was found upon coexpression of the mTalin-YFP marker with AtPIP5K2-mCherry, indicating a stabilizing effect of AtPIP5K2-EYFP, but none of the other fusions tested, on actin dynamics. The asterisk indicates a significant difference from the control (mTalin-YFP/mCherry, non-growing) according to a Student’s T-test (*, p≤0.05).

The data indicate a significantly increased actin occupancy in the pollen tube tip of cells displaying tip swelling upon overexpression of AtPIP5K2-EYFP. The data obtained with 2swap6-EAFP and 6swap2-EYFP indicate the manifestation of intermediate actin distributions when compared to the effects of the parental enzymes, AtPIP5K2 and NtPIP5K6, and correspond well with the intermediate morphological phenotypes observed upon expression of these chimeric variants (see Fig. 1). This observation suggests that, despite their significant effects, the Lin-domains swapped in the chimeras are likely not the only determinants of functional specification of the PI4P 5-kinases tested.

Next, the dynamic motility of actin filaments was assessed by observing growing and non-growing pollen tubes coexpressing different PI4P 5-kinase variants with the live actin marker mTalin-YFP (Fig. 7D, E; Supplemental Movies 9-13). Actin dynamics were recorded over a period of five min by time-resolved SD imaging of z-stacks and subsequent kymographic analysis of maximum intensity projections (Fig. 7D). The motility of actin traces was then quantified according to the kymographs (Fig. 7E). Growing and non-growing pollen tubes coexpressing mTalin-YFP with an mCherry control displayed highly motile actin filaments that were mostly excluded from the pollen tube tip (Fig. 7D, as indicated). Motility of the actin structures is indicated in the kymographs by the absence of horizontal lines. By contrast, actin fibres decorated by mTalin-YFP in cells overexpressing AtPIP5K2-mCherry were substantially stabilized, as evident from the thickness of the observed filaments as well as from the presence of horizontal lines in the kymographic analysis (Fig. 7D, as indicated). Please note that despite the substantial stabilization of actin upon overexpression of AtPIP5K2-mCherry there was still an abundance of highly motile actin filaments (Fig. 7E), indicating that the actin cytoskeleton was still largely functional. The significantly reduced motility of actin filaments was observed in a minor proportion of filaments in cells overexpressing AtPIP5K2-EYFP (Fig. 7E, inset). Please also note that the overexpression of AtPIP5K2-EYFP at a level required to analyze effects associated with tip swelling precludes the observation of growing pollen tubes. When NtPIP5K6-mCherry and mTalin-YFP were coexpressed in pollen tubes, the actin marker again displayed highly motile actin filaments (Fig. 7D, as indicated), as indicated in the kymographs again by the absence of horizontal lines.

The quantitative analysis of actin motility indicates that the expression of AtPIP5K2-mCherry resulted in a significant stabilization of prominent apical actin filaments, concomitant with substantial remaining motility of actin filaments of smaller scale.

### Coordinated localization, interaction and functional interplay of AtPIP5K2 with the actin regulator NtRac5

Based on effects of AtPIP5K2 on actin dynamics, we next aimed to elucidate the molecular basis of PtdIns(4,5)P_2_-dependent regulation of the actin cytoskeleton. Among the known regulators of plant actin dynamics, monomeric GTPases of the Rho of plants (ROP) family are among the best characterized (Kost et al., 1998; Fu et al., 2001; Yalovsky et al., 2008; Poraty-Gavra et al., 2013; Bloch et al., 2016). For the tobacco ROP NtRac5 a link to PtdIns(4,5)P_2_-dependent pollen tube tip swelling has previously been suggested (Kost, 2008; Ischebeck et al., 2011). Importantly, localization of the closely related Arabidopsis ROP6 in plasma membrane nanodomains has recently been demonstrated. Therefore, we next addressed effects of AtPIP5K2-EYFP to ROP signaling (Fig. 8). First, the plasma membrane distribution of AtPIP5K2-EYFP and a coexpressed NtRac5-RFP marker was analyzed by SD (Fig. 8A-D). The maximum intensity projection of confocal z-stacks indicated a general enhanced apical localization of both markers in the pollen tube tip (Fig. 8A), consistent with distribution patterns previously reported for either marker alone (Klahre et al., 2006; Klahre and Kost, 2006; Yalovsky et al., 2008; Stenzel et al., 2012; Grebnev et al., 2017). At a smaller scale, the detailed analysis of the distribution at the membrane surface indicated membrane nanodomains for both AtPIP5K2-EYFP and NtRac5-RFP (Fig. 8B). Plasma membrane nanodomains appeared to cluster into larger islands colabelled by both markers (Fig. 8A,B). The density of the nanodomain signals at the plasma membrane precluded meaningful evaluation of colocalization patterns, which might arise as a consequence of random spatial coincidence. Therefore, we resorted to the quantitative analysis of multiple imaging stacks of lesser complexity as shown in Fig. 8C, where high intensity signals displayed similar localization between AtPIP5K2-EYFP and NtRac5-RFP (boxes in Fig. 8C). Fluorescence signals of multiple partial z-stacks around the median confocal section were analyzed for the sum intensity of the vertical pixel columns (for details, please see the Material and Methods section and Supplemental Fig. S4). The resulting sum intensity profiles for AtPIP5K2-EYFP and for NtRac5-RFP displayed similar patterns, shown for a representative pollen tube in Fig. 8C,D (Fig. 8D upper panel, full range; lower panel, detail), and a tight correlation of fluorescence intensity patterns with a high Pearson coefficient of R=0.95 (Fig. 8E, left panel). The analysis of pollen tube cells coexpressing both markers (N=11 cells, each with five individual measurements) indicates a close correlation of the markers with a mean R-value of 0.87 (Fig. 8E, right panel).

**Figure 8.**
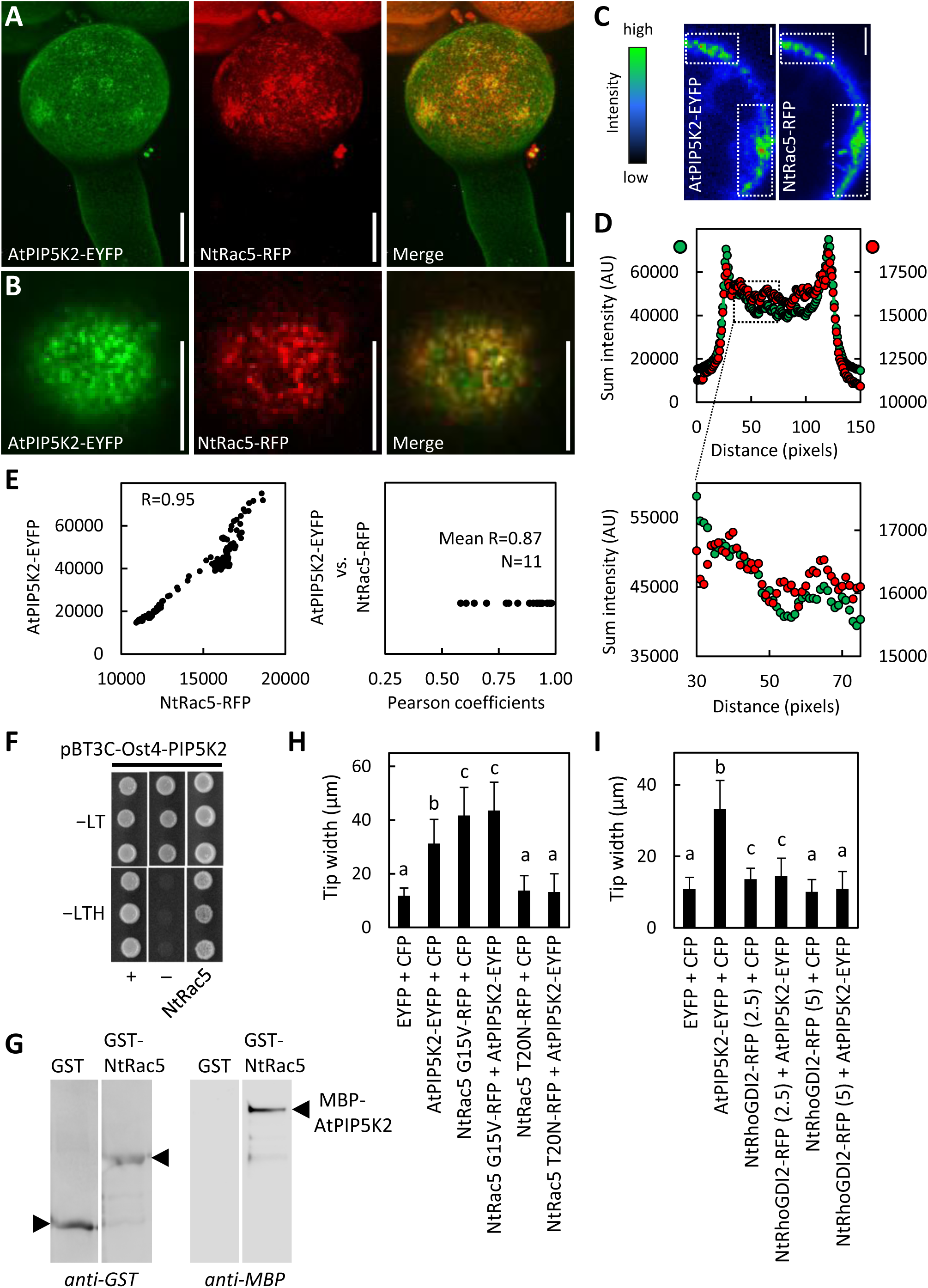
Coordinated localization, interaction and functional interplay of AtPIP5K2 with the actin regulator NtRac5. To explore the molecular mechanicm by which AtPIP5K2 might influence actin dynamics, we tested for links to elements of ROP signalling, which are known to influence pollen tube actin. AtPIP5K2-YFP was coexpressed with the tobacco ROP, NtRAC5-RFP, and fluorescence distributions were monitored by SD. **A**, Representative 3D reconstruction from z-stack acquisition of a pollen tube coexpressing AtPIP5K2-YFP and NtRAC5-RFP, as indicated. Scale bars, 10 µm. **B**, Fluorescence distribution at the cell surface of the pollen tube shown in (A), as indicated. Note that plasma membrane nanodomains of both AtPIP5K2-EYFP and NtRAC5-RFP are visible as bright dots at the cell surface. Scale bars, 10 µm. **C**, Representative median confocal frames extracted from the z-stack projection shown in (A) and representing the individual channels for AtPIP5K2-EYFP or NtRac5-RFP with heatmap colors, as indicated. Note that high intensity values from both channels localize in the same areas at the cell periphery. **D**, The sum intensities were determined for the vertical pixel columns of each channel from regions of interest marked in panel (C) (please refer to Supplementary Fig. S4 and Materials and Methods for details). Upper panel, Sum intensities for AtPIP5K2-YFP (green) and NtRac5-RFP (red). Lower panel, Detail view if the region of interest indicated by the box in the upper panel. **E**, Correlation analysis of sum intensity profiles obtained for AtPIP5K2-EYFP and NtRac5-RFP. Left panel, Representative correlation plot for the fluorescence distributions given in panel (D). The correlation is characterized by a high Pearson coefficient R of 0.95. Right panel, Pearson coefficients obtained accordingly from 11 correlation analyses, with a mean R of 0.87, indicating consistently high correlations between the spatial distributions of AtPIP5K2-EYFP and NtRac5-RFP. **F, G**, AtPIP5K2 and NtRac5 proteins were tested for interaction in a protein complex. **F**, Interaction of AtPIP5K2 and NtRac5 according to split-ubiquitin-based yeast-two-hybrid analysis. In this system, AtPIP5K2 is expressed as a fusion to an Ost4-membrane anchor, which attaches the fusion protein to the cytoplasmic face of the endoplasmatic reticulum. Positive interaction of the protein partners is indicated by yeast growth on selective −LTH media compared to growth of the positive (+) and negative controls (−). Growth of all colonies on −LT media indicates the presence of the correct genotypes of the treansfoprmed yeast tested. The data are shown in triplicates and are representative for 4 biological experiments with similar results. **G**, Interaction of AtPIP5K2 and NtRac5 according to immuno pull-down experiments. The AtPIP5K2 and NtRac5 proteins were recombinantly expressed in E. coli as fusions to N-terminal GST or MBP tags, respectively. The GST-NtRac5 and MBP-AtPIP5K2 proteins were affinity-purified and used for the pull-down experiment. Purified GST protein alone served as a negative control. Positive interaction is indicated by the detection of full-length MBP-AtPIP5K2 upon immunoprecipitation with GST-NtRac5, but not with GST alone. The blots shown are representative for two experiments. **H, I**, Pollen tube tip swelling was used as a morphological marker to assess the functional interplay of AtPIP5K2 with elements of tobacco ROP signalling in vivo. **H**, Functional interplay between AtPIP5K2 and NtRac5. Pollen tube tip widths were determined upon coexpression of the control proteins, CFP with EYFP; AtPIP5K2-EYFP with CFP; the constitutive active NtRac5 G15V-RFP with CFP; AtPIP5K2-EYFP with NtRac5 G15V-RFP; the dominant negative NtRac5 T20N-RFP with CFP; and AtPIP5K2-EYFP with NtRac5 T20N-RFP. Please note the absence of pollen tube tip swelling upon coexpression of AtPIP5K2-EYFP together with NtRac5 T20N-RFP, indicating that activation of NtRac5 is required for the effect of AtPIP5K2-EYFP on tip swelling. Data are given as means ± standard deviation and represent measurements of ≥100 transgenic pollen tubes. **I**, Functional interplay between AtPIP5K2 and NtRhoGDI2. Pollen tube tip widths were determined upon coexpression of the control proteins, CFP with EYFP; AtPIP5K2-EYFP with CFP; NtRhoGDI2-RFP with CFP; and AtPIP5K2-EYFP with NtRhoGDI2-RFP. Numbers indicate the use of 2.5 µg or of 5 µg plasmid DNA for the biolistic transformation of pollen tubes to modulate the expression levels of NtRhoGDI2-RFP. Please note the reduced pollen tube tip swelling upon coexpression of AtPIP5K2-EYFP together with increasing levels of NtRhoGDI2, supporting the notion that AtPIP5K2-EYFP serves as a GDF to mediate pollen tube tip swelling. Data are given as means ± standard deviation and represent measurements of ≥100 transgenic pollen tubes. Letters indicate significan differences according to a one-way Anova test with Tukey post-hoc analysis.

As AtPIP5K2-EYFP and NtRac5-RFP displayed spatially correlated distribution patterns, we next tested for interaction between the AtPIP5K2 and NtRac5 proteins (Fig. 8F, G). An interaction of AtPIP5K2 and NtRac5 in a protein complex is supported by split-ubiquitin-based yeast-two-hybrid analysis, in which cell growth on the restrictive −LTH media indicates positive interaction (Fig. 8F), as assessed compared to the growth of positive (+) and negative controls (−). The interaction was verified by an immuno pull down experiment (Fig. 8G). Both proteins were recombinantly expressed in *E. coli*, AtPIP5K2 as a fusion to an N-terminal maltose-binding protein (MBP) tag (MBP-AtPIP5K2); and NtRac5 as a fusion to an N-terminal glutathione S-transferase (GST) tag (GST-NtRac5). The GST protein was expressed as a negative control and all proteins were affinity purified. Upon coincubation of either GST or GST-NtRac5 with MBP-AtPIP5K2, the MBP-AtPIP5K2 protein was only detected in the immunoprecipitate with GST-NtRac5, not in that with GST alone (Fig. 8G). The immuno-pull down indicates that MBP-AtPIP5K2 bound to GST-NtRac5 but not to GST (Fig. 8G), suggesting physical interaction between the AtPIP5K2 and NtRac5 proteins.

A functional link between AtPIP5K2 and NtRac5 was addressed by monitoring pollen tube tip swelling upon individual or combined expression of these proteins or of relevant protein variants (Fig. 8H). Coexpression of AtPIP5K2-EYFP with a CFP control in pollen tubes resulted in significant tip swelling, compared to control cells coexpressing CFP and EYFP, as indicated (Fig. 8H). Similarly, the coexpression of a constitutive active (CA) variant of NtRac5, NtRac5 G15V-mCherry, with a CFP control also resulted in significant pollen tube tip swelling, as did the coexpression of AtPIP5K2-EYFP with NtRac5 G15V-mCherry, as indicated (Fig. 8H). By contrast, no tip swelling was observed when a dominant negative (DN) variant of NtRac5, NtRac5 T20N-mCherry was expressed. Importantly, the expression of the DN NtRac5 T20N-mCherry abolished the effect of coexpressed AtPIP5K2-EYFP on pollen tube tip swelling (Fig. 8H), indicating that activation of NtRac5 was required for the effect of AtPIP5K2 on pollen tube tip swelling. We conclude that NtRac5 and AtPIP5K2 act in a common pathway to mediate effects on actin and cause pollen tube tip swelling.

To further substantiate this important finding, we next used the coexpression approach to test for functional interplay of AtPIP5K2 with another element of ROP signaling, the pollen expressed tobacco guanidine nucleotide dissociation inhibitor, NtRhoGDI2 (Klahre et al., 2006; Ischebeck et al., 2011; Sun et al., 2015). NtRhoGDI2 can bind NtRac5 in the cytosol, thereby controlling the pool of NtRac5 that can be activated at the plasma membrane (Klahre et al., 2006; Sun et al., 2015). In previous work, it was proposed that PtdIns(4,5)P_2_ can serve as a GDI-displacement factor (GDF) (Faure et al., 1999) and that in pollen tubes PtdIns(4,5)P_2_ may promote the release of NtRac5 from the NtGHDI2/NtRac5 complex (Kost, 2008; Ischebeck et al., 2011). AtPIP5K2-mediated pollen tube tip swelling might thus be explained at the molecular level by enhanced PtdIns(4,5)P_2_ production promoting the release and activation of NtRac5 at the plasma membrane, with ensuing stabilizing effects on actin and tip swelling (Klahre et al., 2006; Ischebeck et al., 2011). When AtPIP5K2-EYFP was coexpressed with a CFP control, we observed significant tip swelling compared to control pollen tubes expressing EYFP and CFP (Fig. 8I, as indicated), as in previous experiments. By contrast, decreasing degrees of pollen tube tip swelling were observed when AtPIP5K2-EYFP was coexpressed with NtRhoGDI2-RFP, using increasing amounts of the *NtRhoGDI2-RFP* cDNA for particle bombardment to achieve increasing expression levels. The data indicate that the co-overexpression of NtRhoGDI2-RFP canceled the effects of AtPIP5K2-EYFP, which is consistent with the previously proposed function of PtdIns(4,5)P_2_ as a GDF.

Together, the data provide evidence that AtPIP5K2 localizes in plasma membrane nanodomains of pollen tubes in close spatial proximity with NtRac5. AtPIP5K2 interacts physically with NtRac5, and both proteins act in a common pathway to effect pollen tube tip swelling, likely through a regulatory effect on pollen tube actin dynamics.

## Discussion

In this study, we addressed subcompartmentalization as an aspect of localized functional specification of the plasma membrane, with a focus on membrane nanomains containing the regulatory phospholipid, PtdIns(4,5)P_2_. For the functional aspects of our study, we exploited the unique properties of the tobacco pollen tube model, where alternative functions of the PI4P 5-kinases AtPIP5K2 and NtPIP5K6 have previously been found to occur within a narrow subapical region of the plasma membrane.

### Nano-organization of PtdIns(4,5)P_2_ in the pollen tube plasma membrane

The subcompartmentalization of plasma membrane proteins in membrane nanodomains - or possibly “rafts” (Simons and Toomre, 2000; Simons and Vaz, 2004) - has previously been proposed for plants based on the subcellular localization of fluorescent markers for proteins identified, e.g., by proteomic analysis of detergent-insoluble membrane (DIM) preparations (Mongrand et al., 2004), such as remorins (Morel et al., 2006). Fluorescent variants of such markers show clear association with membrane nanodomains when analyzed microscopically and have been used as markers for nanodomain structure and membrane organization in the context of physiological processes mostly related to pathogen infection (Lefebvre et al., 2007; Raffaele et al., 2007; Raffaele et al., 2009; Lefebvre et al., 2010; Jarsch and Ott, 2011; Marin and Ott, 2012; Toth et al., 2012; Bozkurt et al., 2014; Jarsch et al., 2014; Konrad et al., 2014; Bucherl et al., 2017; Liang et al., 2018). While previous studies have focused mainly on the nanodomain association of proteins, it is evident that membrane lipids will also be essential determinants of membrane nanodomain formation. Based on our SD imaging of the distribution of the PtdIns(4,5)P_2_ biosensor, RedStar-PLC-PH, in the pollen tube plasma membrane, we observed the subcompartmentalization of the biosensor into nanodomains (Fig. 2). This pattern is consistent with previous reports, for instance because PtdIns(4,5)P_2_ has been proposed to be essential for the membrane insertion of nanodomain-associated remorins (Gronnier et al., 2017). The nano-organization of PtdIns(4,5)P_2_ in the pollen tube plasma membrane provides a rationale for coexisting alternative roles of PtdIns(4,5)P_2_.

### PtdIns(4,5)P_2_-nanodomains influence ROP signalling

The comparison of the functionally divergent PI4P 5-kinases AtPIP5K2 and NtPIP5K6 suggests that PtdIns(4,5)P_2_ formed in nanodomains exerts a different effect than PtdIns(4,5)P_2_ outside the domains (see Figs. 1, 3). Specifically, actin stabilization and pollen tube tip swelling emerged as a consequence of PtdIns(4,5)P_2_ formed in plasma membrane nanodomains defined by AtPIP5K2. PtdIns(4,5)P_2_ has previously been proposed to mediate pollen tube tip swelling through effects on the actin cytoskeleton and by circumstantial evidence been linked to ROP signalling (Ischebeck et al., 2011). This notion is of particular interest, because Arabidopsis ROP6 has recently been found to associate with dynamic plasma membrane nanodomains (Platre et al., 2019). These ROP6 nanodomains were proposed to be sensitive to the abundance of phosphatidylserine (Platre et al., 2019), leading us to hypothesize that also other anionic lipids, such as PtdIns(4,5)P_2_, might contribute to the nano-organization of ROPs. The effects of overexpressed AtPIP5K2-EYFP on in vivo actin dynamics in pollen tubes (Fig. 7) and the coordinated localization and physical interaction of AtPIP5K2 and the ROP, NtRac5 (Fig. 8), are consistent with an effect of PtdIns(4,5)P_2_ on ROP signalling and cytoskeletal control. More specifically, the functional interplay of AtPIP5K2 with NtRac5 (Fig. 8H) and the guanidine nucleotide dissociation inhibitor (GDI) NtRhoGDI2 (Fig. 8I) provides experimental evidence that PtdIns(4,5)P_2_ acts as a GDI displacement factor (GDF), as previously proposed (Kost, 2008), enabling the enhanced release of NtRac5 to the plasma membrane for activation (Faure et al., 1999; Kost, 2008; Ischebeck et al., 2011).

### PtdIns(4,5)P_2_ and phosphatidylserine may influence different aspects of plasma membrane nano-organization

The nano-organization of the plasma membrane may enable interactions of PtdIns(4,5)P_2_ or its downstream effectors, such as NtRac5, with specific partners, thus enabling divergent functional consequences, as outlined in the simplified model scheme (Fig. 9). It appears plausible that plasma membrane nanodomains serve as platforms for proteins, such as AtPIP5K2 and NtRac5, which assemble into functional complexes to influence the membrane attachment and stabilization of actin filaments (Fig. 9A). It is possible that the localized attachment of actin filaments to the plasma membrane also serves as an initiating factor for the assembly of proteins in membrane nanodomains, or that both nanodomains and actin exert mutual effects in a self-organizing manner. Within plasma membrane nanodomains, PtdIns(4,5)P_2_ might influence NtRac5 availability for activation (Fig. 9B), whereas - in analogy to the situation reported for ROP6 in Arabidopsis (Platre et al., 2019) - phosphatidylserine possibly might influence lateral distribution of NtRac5. ROPs exert their regulatory effects by signaling through Rho-interactive CRIB-domain containing (RIC) proteins (Gu et al., 2005; Lee et al., 2008). In particular, RIC proteins have been proposed to mediate the coordination of actin dynamics and secretion in pollen tubes (Lee et al., 2008), our data suggest that the overproduction of PtdIns(4,5)P_2_ in membrane nanodomains may shift NtRac5 effects towards RIC4-dependend F-actin accumulation, whereas overproduction of PtdIns(4,5)P_2_ outside of membrane nanodomains might favor RIC3-dependent depolymerization of F-actin and promote exocytosis. In this scenario, ensuing overactivation of NtRac5 might result in the stabilization of cortical actin in the pollen tube tip through tobacco RIC4 homologs, possibly resulting in reduced rates of secretion and vesicle accumulation (Lee et al., 2008). PtdIns(4,5)P_2_ within or outside of membrane nanodomains may thus serve to direct NtRac5 towards different RICs, thereby aiding the coordination between actin dynamics and secretion during pollen tube growth. The combined data on plasma membrane nanodomains for PtdIns(4,5)P_2_ and AtPIP5K2 indicate that spatial separation of PtdIns(4,5)P_2_ populations is one reason for the functional distinction previously reported for AtPIP5K2 vs. other PI4P 5-kinases of subfamily B (Ischebeck et al., 2008; Ischebeck et al., 2010b; Stenzel et al., 2012).

**Figure 9.**
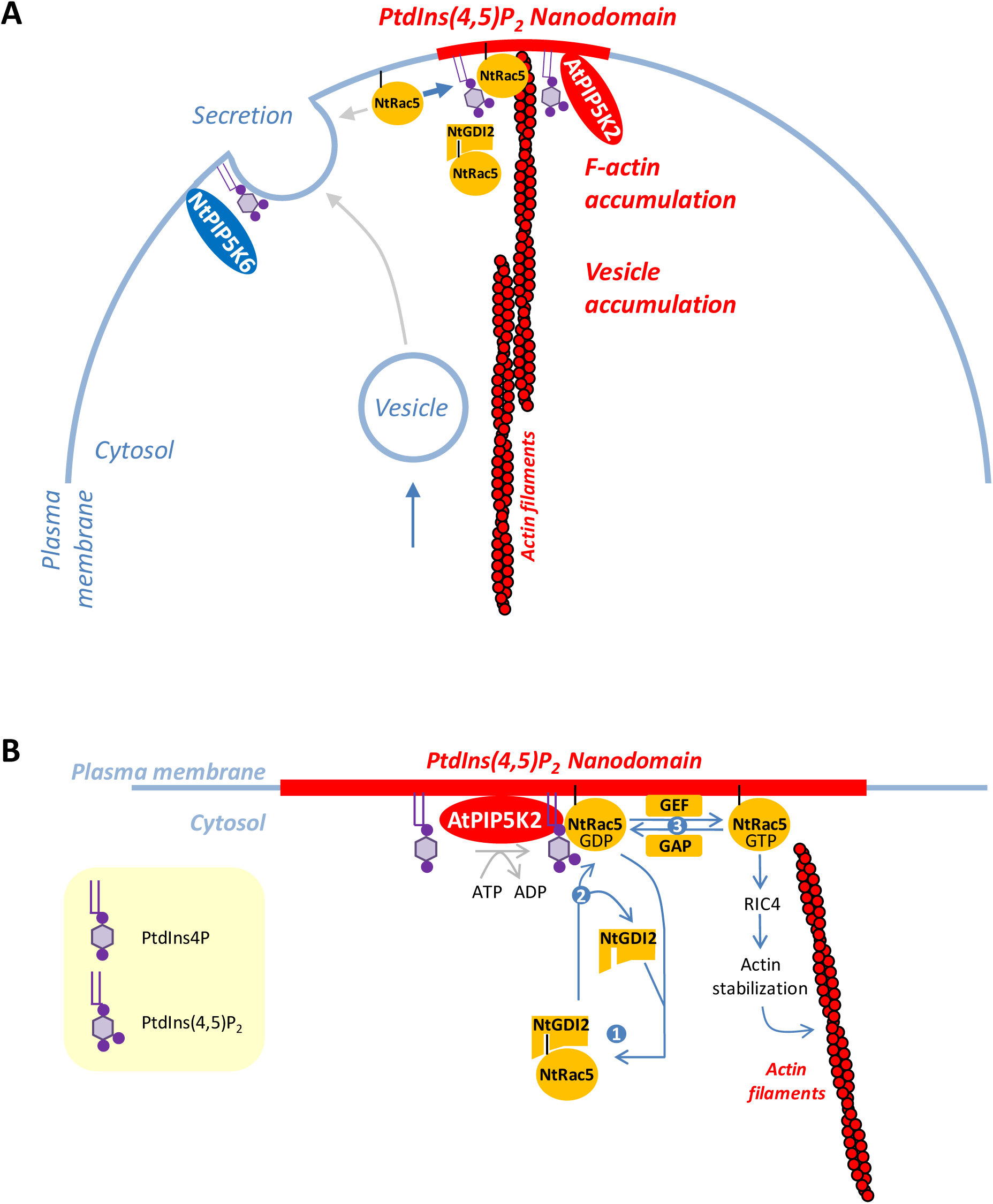
Model for the influence of nanodomain-associated PtdIns(4,5)P_2_ and AtPIP5K2 on NtRac5 and actin dynamics. Polar tip growth of pollen tubes requires the close spatio-temporal coordination of actin dynamics and secretion/endocytosis. **A**, Nano-organization of the plasma membrane with AtPIP5K2 associating with nanodomains (red) and NtPIP5K6 distributing uniformly (blue) in the plasma membrane. PtdIns(4,5)P_2_ formed by AtPIP5K2 in nanodomains may act to stabilize the actin cytoskeleton (red filaments) through functional interplay with elements of ROP signalling, such as NtRac5 or NtRhoGDI2 (NtRhoGDI2). Upon overproduction of PtdIns(4,5)P_2_ in nanodomains, such as upon overexpression of AtPIP5K2, NtRac5-dependent effects might be shifted towards promoting F-actin accumulation (thick blue arrow), possibly at the expense of other functions (thin grey arrows). The diffusely localized PtdIns(4,5)P_2_ formed by NtPIP5K6 serves a role in promoting secretion, as indicated. **B**, Proposed functional interplay of nanodomain-associated PtdIns(4,5)P_2_ with ROP signalling. PtdIns(4,5)P_2_ may act as a GDF on NtRhoGDI2 to promote NtRac5-dependent stabilization of the actin cytoskeleton. A pool of plasma membrane-associated NtRac5 is in a dynamic equilibrium with cytosolic NtRac5 bound to NtRhoGDI2 (point 1). PtdIns(4,5)P_2_ formed by AtPIP5K2 in plasma membrane nanodomains might act as a GDF to destabilize cytosolic NtRhoGDI2/NtRac5 complexes (point 2), thereby releasing more NtRac5 to the plasma membrane. At the plasma membrane, NtRac5 can be activated by GEFs to NtRac5-GTP, which initiates a signalling cascade to stabilize actin (point 3). Other explanations are possible. GAP, GTPase-activating protein; GEF, Guanidine nucleotide exchange factor; PtdIns4P, phosphatidylinositol 4-phosphate; PtdIns(4,5)P_2_, phosphatidylinositol 4,5-bisphosphate; RIC, ROP-interactive CRIB-domain containing protein

### Plasma membrane nano-organization and limited lateral diffusion of PtdIns(4,5)P_2_

It is important to note that the observed spatial distribution patterns of PtdIns(4,5)P_2_ or PI4P 5-kinase isoforms can help to explain functional consequences of PtdIns(4,5)P_2_-dependent processes for pollen tube growth only if we assume that spatially separated PtdIns(4,5)P_2_ pools do not mix before their respective downstream effects have taken place. Thus, the observed patterns lead us to postulate limited lateral diffusion of PtdIns(4,5)P_2_ molecules from their sites of biosynthesis, which are defined by the localization of the different PI4P 5-kinase isoforms, i.e., AtPIP5K2 in nanodomains vs. NtPIP5K6 or AtPIP5K5 in continuous membrane patterns. While this concept may oversimplify, it is consistent with current models of membrane organization and restricted lateral movement of membrane components (Arumugam et al., 2015; Destainville et al., 2016). Current models on the structure of membrane nanodomains are mostly based on the raft hypothesis (Rajendran and Simons, 2005) and assume that a complex mixture of membrane lipids, including sphingolipids, sterols and phosphoinositides, may form a joint structure with specific biophysical properties (Anishkin and Kung, 2013), facilitating certain membrane functions while excluding others. For instance, attachment of cytoskeletal strands to the plasma membrane may require a localized rigidification around the sites of anchorage (Anishkin and Kung, 2013). Nanodomain organization may also create biophysical barriers for the lateral diffusion of intrinsic membrane proteins as well as for membrane lipids, such as PtdIns(4,5)P_2_ (Gerth et al., 2017). Considering the raft hypothesis (Simons and Ikonen, 1997; Simons and Toomre, 2000; Simons and Vaz, 2004), the finding of AtPIP5K2 and a PtdIns(4,5)P_2_ reporter in plasma membrane nanodomains is consistent with the previous report of PI4P 5-kinase activity associated with detergent-resistant membrane (DIM) fractions prepared from plasma membrane from tobacco leaves (Furt et al., 2010). Of note, PI4P 5-kinase activity was previously reported to reside in DIM preparations as well as outside of DIMs of tobacco leaves or bright yellow 2 (BY-2) cells (Furt et al., 2010), supporting our data indicating PtdIns(4,5)P_2_ production within and outside of plasma membrane nanodomains (Figs. 2 and 4). Formation of PtdIns(4,5)P_2_ by different PI4P 5-kinase isoforms, such as AtPIP5K2 within or NtPIP5K6 outside of nanodomains may determine the downstream effects of PtdIns(4,5)P_2_ possibly through binding to alternative protein partners, resulting in the control of different PtdIns(4,5)P_2_-dependent processes (Fig. 9A). Localization of the enzymes in membrane nanodomains may thus facilitate the interaction with partners involved in the control of the action cytoskeleton, such as ROPs (Fig. 9B, C), and enable differential regulation of PtdIns(4,5)P_2_ effects in different membrane areas.

While our data suggest a distinct regulatory function of PtdIns(4,5)P_2_ in plasma membrane nanodomains, it appears likely that the “diffuse” PtdIns(4,5)P_2_ outside of the nanodomains may also be further subcompartmentalized. Besides a distinction between PtdIns(4,5)P_2_ pools based on spatial separation, which may or may not be detectable by microscopy, it is possible that diffusely localizing PI4P 5-kinases may display a patchy pattern of activated and inactivated states, resulting in islands of active PI4P 5-kinases within a population of possibly not activated proteins that is diffusely distributed. This hypothesis is based on recent reports that the type B PI4P 5-kinase AtPIP5K6 is regulated in its activity by the mitogen-activated protein kinase (MAPK) MPK6, and that MPK6-mediated phosphorylation does not impair membrane association of PIP5K6 (Hempel et al., 2017; Menzel et al., 2019).

### PtdIns(4,5)P_2_ nanodomains only in pollen tubes and other open questions

The pollen tube system used in this study holds several unique advantages, including high levels of PtdIns(4,5)P_2_ (Supplemental Fig. S5 A) and the previous characterization of divergent functional effects of AtPIP5K2 and NtPIP5K6 on pollen tube morphology. In previous studies, a contribution of phosphoinositides to the regulation of important physiological processes has also been reported for the vegetative portion of plants. These studies describe roles for phosphoinositides in clathrin recruitment (König et al., 2008; Ischebeck et al., 2013); the polarization of auxin efflux carriers of the PIN-FORMED (PIN)-family (Mei et al., 2012; Ischebeck et al., 2013; Tejos et al., 2014), of D6P-kinases (Barbosa et al., 2016) and of BRX and PAX proteins (Marhava et al., 2020); the control of cytokinesis (Lin et al., 2019); cell wall deposition (Rodriguez-Villalon et al., 2015), and roles in membrane trafficking during plant pathogen interactions (Shimada et al., 2019; Qin et al., 2020). While these reports together may suggest the presence of PtdIns(4,5)P_2_ in membrane nanodomains occupied by phosphoinositides, in Arabidopsis hypocotyl cells we detected such nanodomains by SD only upon application of salt (Supplemental Fig. S5 B), thus artificially increasing PtdIns(4,5)P_2_ levels. As a chemical difference has previously been found between PtdIns(4,5)P_2_ molecular species in resting and salt-stressed Arabidopsis plants (König et al., 2007; König et al., 2008), the nature of the punctate signals seen in the hypocotyl cells only upon salt stress remains unclear. Therefore, it remains unclear whether our data from the gametophytic pollen tube system can be transferred to sporophytic cells of the plant.

Our study also does not address a contribution of other lipid classes to the formation of PtdIns(4,5)P_2_-containing plasma membrane foci, so it remains open, whether or not the lipid raft concept may be applied to the nanodomains observed here. The previous observations that defects in the biosynthesis of sterols (Men et al., 2008), sphingolipids (Markham et al., 2011) and phosphoinositides (Mei et al., 2012; Ischebeck et al., 2013; Tejos et al., 2014) all result in similar mistrafficking of PIN proteins, are in favour of the notion that these lipids form a joint functional membrane structure underlying these effects in sporophytic tissues. An oxysterol-binding protein fused to a fluorescent reporter has previously been shown to decorate a patchy pattern in the subapical plasma membrane of pollen tubes (Skirpan et al., 2006), suggesting the presence of sterol domains in the plasma membrane of pollen tubes. Furthermore, oxysterol-binding proteins from the yeast *Saccharomyces cerevisiae* contain PH-domains which specifically bind phosphoinositides (Roy and Levine, 2004; Fairn et al., 2007) and influence Rho-dependent cell polarization (Kozminski et al., 2006). In the absence of further direct evidence, however, the contribution of other lipid classes to the formation of membrane nanodomains enriched in PtdIns(4,5)P_2_ remains to be analyzed. Future experiments will be directed towards further unraveling the nano-organization and chemical composition of plant plasma membrane nanodomains.

## Methods

### cDNA constructs

The assembly of plasmids encoding AtPIP5K2-EYFP, AtPIP5K2-mCherry, NtPIP5K6-EYFP, NtPIP5K6-mCherry, AtPIP5K2_344-754_-EYFP, 2_swap6-EYFP and 6swap2-EYFP was previously described (Stenzel et al., 2012). The RedStar-PLC-PH construct was assembled as previously described (Ischebeck et al., 2010b). The Lifeact-RFP cassette (Berepiki et al., 2010) in pAL10 vector, kindly provided by Dr. Stephan Seiler (University of Freiburg, Germany), was digested with SalІ and NotІ and transferred into the pEntryD vector. The tomato Lat52 promoter was amplified with Sfi1 sites from the pLATGW (Twell et al., 1990) and placed in front of the Lifeact-RFP cassette. For Lifeact-EYFP, RFP was then replaced with EYFP. The mTalin-YFP construct was a kind gift of Prof. Dr. Benedikt Kost (University of Erlangen, Germany). For yeast two hybrid analysis, the cDNA of the *At*PIP5K2 bait protein was cloned into the plasmid *pBT3-C-OST4*. *pBT3-C-OST4* was produced by inserting the cDNA of OST4 (*yeast oligosaccharyl transferase 4*) into the *Xba*I site upstream of the mcs of the original pBT3-C plasmid (*Dualsystems* Biotech AG, Zürich, Switzerland) as previously described (Möckli et al., 2007). The At*PIP5K2* cDNA was amplified with the primer combination At*PIP5K2-OST4-for*/At*PIP5K2-OST4-rev*.

The primers introduced an *Sfi*IA site upstream and an *Sfi*IB site downstream of the At*PIP5K2* sequence, and the PCR fragment was inserted directionally into the *Sfi*I sites of the *pBT3-C-OST4* vector. The cDNAs for the prey protein NtRac5 was amplified by *Rac5-SfiIA.for*/ *Rac5-SfiIB.rev* and moved as *Sfi*I fragments into the mcs of pPR3-N (*Dualsystems* Biotech AG, Zürich, Switzerland).

### Transient expression of constructs in tobacco pollen tubes

Mature pollen grains were collected from four to five tobacco flowers of approximately eight to nine-week-old plants. Pollen were resuspended in liquid pollen tube growth media (Read et al., 1993) followed by the filtering of the pollen grains onto a cellulose acetate filter, and then transferred into Whatmann paper which was moistened with pollen tube growth media. They were immediately transformed by bombardment with plasmid-coated 1-μm gold particles with a helium-driven particle accelerator (PDS-1000/He; Bio-Rad) using 1350 psi rupture disks and a vacuum of 28 inches of mercury. Prior to the bombardment, these gold particles were coated with 4-5 µg of the desired plasmid of interest. After bombardment, the pollen grains were transferred into 300 µl of pollen tube growth media which was then equally divided onto three microscopic glass slides and viewed under the microscope 4-6 hours after the bombardment.

### Live cell microscopy and image processing

Images were acquired with a Zeiss LSM880 Airyscan system or with a Zeiss Cell observer SD with a Yokogawa CSU-X1 spinning disc unit. For imaging with the LSM880, a 20x or a 63x oil immersion objective was used and images were captured with the ZEN Black image analysis software. During acquisition with the LSM880, EYFP was excited at 514 nm laser line and imaged with a HFT 514 nm major beam splitter (MBS) while mCherry or RFP was excited with a 561nm laser line and imaged with a HFT 561nm MBS. The Zeiss Cell observer SD was used for acquiring time – lapse images of pollen tubes expressing YFP or mCherry fluorophores which was acquired with the Zen Blue image analysis software. Images were acquired with a 100x oil immersion objective and captured with a Photometrics Evolve 512 Delta EM-CCD camera. YFP was excited at 491 nm and mCherry or RedStar at 561 nm using a multichannel dichroic and an ET525/50M or an ET595/50M band pass emission filter (Chroma Technology) for GFP and mCherry respectively.

All images were processed using Fiji software (Schindelin et al., 2012). Nanodomain lifetimes were first acquired using the SD microscope where images were captured every two s over a period of 120 s. The images were then processed using the Fiji ‘Trackmate’ plugin (Tinevez et al., 2017). With the same plugin the x;y coordinates of nanodomains were extracted and the Euclidean distances from the tip measured (x;y coordinates of each pollen tube tip analyzed were manually extracted). For size determination of PtdIns(4,5)P_2_ nanodomains, punctate signals were manually picked and listed into a ROI manager. To assess the intensity distributions of AtPIP5K2-EYFP, NtPIP5K6-EYFP, 2swap6-EYFP and 6swap2-EYFP at the plasma membrane, the Fiji software was used. An area of 22×22 pixels at the cell surface of multiple pollen tubes was manually picked, listed into the ROI manager and the standard deviation among pixel values was calculated. High standard deviations reflect the presence of heterogeneous intensity distribution at the cell membrane, and therefore foci formation; by contrast, low standard deviations reflect a uniform distribution of intensoties, and therefore diffuse distribution of the marker.

For correlation analysis of AtPIP5K2-EYFP with RedStar-PLC-PH and of AtPIP5K2-EYXFP with NtRac5-RFP, multiple pollen tubes were analysed, each with five z-stacks. For each image, fluorescence channels (green and red) were split and individually analysed. Pixel intensities were extracted using Fiji; every image was then divided into longitudinal sections of 1 pixel width, the intensity values of each pixel column was summed up and plotted as a function of space. The product momentum Pearson correlation coefficient (R) among the two channels was then determined (please also look at Supplementary Fig. S4).

Actin occupancy was determined by using the pixel classification and object classification tool from the machine-learning software Ilastik (Berg et al., 2019); pixel occupancy of lifeact fluorescence signal of each pollen tube was extracted, divided by the area of the pollen tube tip and values were converted into area percentages. To determine the ratio of plasma membrane vs. cytosolic fluorescences, the same software tool was used to extract the signal intensities from imaging frames.

Actin dynamics were monitored over time in full z-stacks from pollen tubes transiently transformed with mTalin-YFP, with image acquisition at every 10 s over a period of 5 min (resulting in 30 frames), using LSM. Kymographs were then extracted for every pollen tube using Fiji software. In particular, to focus on actin dynamics in the pollen tube tip, for every image an area of 28×25 micron at the tip was cropped and analyzed. Kymograph data was recorded at the center of the tube. Dynamic actin structures are apparent by this analysis as short and/or asymmetric fluorescent signals in the kymograph. By contrast, stabilized/static actin structures appear in the kymographs as long fluorescent lines. Fluorescence signals from each kymograph were segmented and analyzed using the pixel classification and object classification tool from Ilastik software (Berg et al., 2019). The lengths of each fluorescence signal were extracted in pixels (Euclidean diameter). Fluorescence signals shorter than 5 pixels were considered as noise and were excluded from measurements.

For pectin staining pollen tubes were stained as previously reported (Iwai et al., 2006; Ischebeck et al., 2008) with ruthenium red (Sigma-Aldrich) using a final concentration of 0.01% (w/v) and imaged under the light microscope (Zeiss Axiovert 200) within 5 to 15 min after addition of the dye.

For FM 4-64 staining (Molecular Imaging Products), the dye was added to pollen tubes to a final concentration of 2.5 µM and with an incubation time of 2-3 minutes before imaging.

### Recombinant protein expression

Extracts containing recombinant protein were generated by expression in *Escherichia coli* Rosetta 2 cells. Cells were grown in liquid LB media selecting for kanamycin and chloramphenicol resistance. Expression cultures of 100 ml were incubated at 37°C shaking at 200 rpm. Expression was induced at an optical density of 0.8 by adding 1 mM isopropyl-β-D-thiogalactosylpyranoside (IPTG). Cultures were incubated over night at 25°C shaking at 200 rpm and harvested by centrifugation. The cell sediment was resuspended on ice in lysis buffer containing 50 mM Tris-HCl, pH 8, 300 mM NaCl, 1 mM EDTA, 10 % (w/v) glycerine and 50 U of lysozyme. Cells were ruptured by ultrasound on ice, using five 60 s bursts in a Sonifier Cell Disruptor B15 (Branson Dietzenbach, Germany) at 50 % power and 50 % impulse settings. The protein extracts were cleared by centrifugation and stored at −20°C.

### Protein interaction studies

The split-ubiquitin-based yeast two-hybrid-system (Dualsystems Biotech, Zürich, Switzerland (Johnsson and Varshavsky, 1994)) was used to analyze protein-protein interactions between NtRac5 and PIP5K2 as bait, which was expressed as a fusion to an N-terminal OST4-anchor (Möckli et al., 2007). All analyses were performed according to the manufacturer’s instructions. For immuno-pull-down experiments, recombinant GST, GST-NtRac5 were immobilized on glutathione-agarose (Thermo Fisher Scientific) and incubated with purified recombinant MBP-AtPIP5K2 protein for 60 min at 4°C. Upon washing of the resin, GST-bound proteins were eluted with 50 mM glutathione. Interacting MBP-AtPIP5K2 protein was detected using an anti-MBP antibody (NEB). Protein input was detected using an anti-GST antibody (GE Healthcare).

### Statistical Evaluation

All quantitative data were tested for statistical significance using two-tailed Student’s t tests. Confidence intervals are given in the figure legends for each data set.

### Accession Numbers

Sequence data from this article can be found in the GenBank/EMBL data libraries under the following accession numbers: AtPIP5K2, At1g77740; AtPIP5K5, At2g41210; NtPIP5K6, JQ219669; NtRac5, AJ250174; NtRhoGDI2, DQ416769.

## Acknowledgements

cDNA clones for mTalin, NtRac5, NtRac5 G15V, NtRac5 T20N and NtRhoGDI2, were kindly provided by Prof. Dr. Benedikt Kost (University of Erlangen, Germany). The cDNA encoding Lifeact was a kind gift of Dr. Stephan Seiler (University of Freiburg, Germany). Financial support by the German Research Foundation (DFG, grants He3424/1-1, He3424/6-1, He3424/6-2, CRC648 TP B10 and INST271/371-1 FUGG to I.H.) is gratefully acknowledged.

## Author Contributions

MF, PK, IS, MH and MR performed experiments; MF, PK, IS, MH, KB, and IH analyzed data; MF, PK and IH designed research; IH wrote the manuscript.

## Legends to Supplemental Figures

**Supplemental Figure S1. Phylogenetic relations of plant PI4P 5-kinases, including AtPIP5K2 and NtPIP5K6 used in this study.** To construct the phylogenetic tree, AtPIP5K amino acid sequences were aligned using the MAFFT multiple sequence alignment program with the L-INS-i option (Katoh et al., 2005). After manual editing of the alignment, ProtTest 3.0 was used for selection of the best-fit model of protein evolution (Abascal et al., 2005). Phylogenetic relationships based on amino acid sequences were inferred from maximum likelihood methods implemented in MEGA 5 (JTT+G+F, 1000 bootstrap replicates) as previously described (Tamura et al., 2011). The phylogenetic tree was mid-point rooted and edited with FigTree v1.3.1 (http://tree.bio.ed.ac.uk/software/figtree).

Supports Fig. 1.

**Supplemental Figure S2. Influence of expressed fluorescence markers on pollen tube growth rates.** The growth rates in µm min^-1^ were determined for untransformed pollen tubes, for pollentubes expressing an EYFP control or for tubes expressing the RedStar-PLC-PH biosensor used to analyze PtdIns(4,5)P_2_ distributions. Data are means ± standard deviation calculated from 30 cells each. The asterisk indicates a significant difference according to a Student’s T-test (*, p≤0.05).

Supports Figs. 2 and 3.

**Supplemental Figure S3. Diffuse localization of AtPIP5K5-EYFP, a PI4P 5-kinase promoting apical pectin secretion in pollen tubes.** The fluorescence distribution of AtPIP5K5-EYFP and AtPIP5K2-EYFP were monitored side-by-side upon transient expression in tobacco pollen tubes by SD. A, Localization of AtPIP5K5-EYFP in the apical plasma membrane (median confocal section). B, Localization of AtPIP5K5-EYFP in the apical plasma membrane, observed at the cell surface (n = 16 transformed pollen tubes from three independent experiments). Dashed line, trace recorded for the intensity profile in (C). C, Fluorescence intensity profile of AtPIP5K5-EYFP. D, Localization of AtPIP5K2-EYFP in the apical plasma membrane (median confocal section). E, Localization of AtPIP5K2-EYFP in the apical plasma membrane, observed at the cell surface (n = 18 transformed pollen tubes from three independent experiments). Dashed line, trace recorded for the intensity profile in (F). F, Fluorescence intensity profile of AtPIP5K2-EYFP.

Supports Fig. 4.

**Supplemental Figure S4. Analytic work flow for the analysis of fluorescence intensity distributions of irregularly-shape structures by extracting vertical pixel columns and determining the sum intensities.** Details as given in the figure.

Supports Figs. 5 and 8.

**Supplemental Figure S5. Analysis of PtdIns(4,5)P_2_ in different plant tissues.**

**A**, The relative levels of PtdIns(4,5)P_2_ were compared based on the biochemical analysis of unlabeled PtdIns(4,5)P_2_ by combined thin-layer chromatography and quantification of the lipid-associated fatty acids by gas chromatography. Control, Whole Arabidopsis seedlings. Please note the high levels of PtdIns(4,5)P_2_ in pollen tubes when compared to vegetative plant tissues.

**B**, The formation of membrane nanodomains was assessed by SD in hypocotyl cells of transgenic Arabidopsis plants expressing the PtdIns(4,5)P_2_ biosensor, YFP_PLCd1-PH_. This biosensor contains the identical lipid-binding region as RedStar-PLC-PH used in this study. No nanodomains decorated by YFP_PLCd1-PH_ were detected in control cells, whereas nanodomains were formed upon application of exogenous salt stress (200 mM). This observation is consistent with low levels of PtdIns(4,5)P_2_ in vegetative tissues of Arabidopsis (see A), and with salt-induced increases in PtdIns(4,5)P_2_ in Arabidopsis. Scale bars, 10 µm. Right panel, Plasma membrane nanodomains observed upon salt-stress did not display detectable dynamics according to kymograph analysis.

Supports Fig. 9.

## Legends to Supplemental Movies

**Supplemental Movie 1. The PtdIns(4,5)P_2_-specific biosensor RedStar-PLC-PH decorates plasma membrane nanodomains in tobacco pollen tubes.** z-stack of confocal sections of 0.9 µm acquired by SD. Supports Fig. 2.

**Supplemental Movie 2. The PtdIns(4,5)P_2_-specific biosensor RedStar-PLC-PH decorates plasma membrane nanodomains in tobacco pollen tubes.** 3D projection of the confocal z-stack shown in Supplemental Movie 1. Supports Fig. 2.

**Supplemental Movie 3. Bright field imaging of a growing tobacco pollen tube transformed with RedStar-PLC-PH.** Images were acquired at 0.1 frames s^-1^. Label, min:s. Supports Fig. 3.

**Supplemental Movie 4. Dynamic fluorescent plasma membrane nanodomains in a growing pollen tobacco pollen tube expressing RedStar-PLC-PH.** Images are from the same pollen tube as shown in Supplemental Movie 3 and were acquired by SD at 0.5 frames s^-1^. The confocal plane is focused on the cell surface. PtdIns(4,5)P_2_ nanodomains are visible as dynamic bright dots. Label, min:s. Supports Fig. 3.

**Supplemental Movie 5. Bright field imaging of a non-growing tobacco pollen tube transformed with RedStar-PLC-PH.** Images were acquired at 0.1 frames s^-1^. Label, min:s. Supports Fig. 3.

**Supplemental Movie 6. Dynamic fluorescent plasma membrane nanodomains in a non-growing pollen tobacco pollen tube expressing RedStar-PLC-PH.** Images are from the same pollen tube as shown in Supplemental Movie 5 and were acquired by SD at 0.5 frames s^-1^. The confocal plane is focused on the cell surface. PtdIns(4,5)P_2_ nanodomains are visible as dynamic bright dots. Label, min:s. Supports Fig. 3.

**Supplemental Movie 7. Bright field imaging of a tobacco pollen tube expressing AtPIP5K2-EYFP.** Images were acquired at 0.1 frames s^-1^. Label, min:s. Supports Fig. 7.

**Supplemental Movie 8. Dynamic fluorescent plasma membrane nanodomains in a tobacco pollen tube expressing AtPIP5K2-EYFP.** Images are from the same pollen tube as shown in Supplemental Movie 7 and were acquired by SD at 0.5 frames s^-1^. The confocal plane is focused on the cell surface. AtPIP5K2-EYFP nanodomains are visible as dynamic bright dots. Label, min:s. Supports Fig. 7.

**Supplemental Movie 9. Actin dynamics in a growing tobacco pollen tube coexpressing mCherry and mTalin-YFP.** z-stacks of 0.9 µm confocal sections through the entire depth of the cell were acquired by LSM at 0.1 frames s^-1^. Only the YFP-channel is shown. Label, min:s. Supports Fig. 7.

**Supplemental Movie 10. Actin dynamics in a non-growing tobacco pollen tube coexpressing mCherry and mTalin-YFP.** z-stacks of 0.9 µm confocal sections through the entire depth of the cell were acquired by LSM at 0.1 frames s^-1^. Only the YFP-channel is shown. Label, min:s. Supports Fig. 7.

**Supplemental Movie 11. Actin dynamics in a pollen tube coexpressing AtPIP5K2-mCherry and mTalin-YFP.** z-stacks of 0.9 µm confocal sections through the entire depth of the cell were acquired by LSM at 0.1 frames s^-1^. Only the YFP-channel is shown. Label, min:s. Supports Fig. 7.

**Supplemental Movie 12. Actin dynamics in a growing pollen tube coexpressing NtPIP5K6-mCherry and mTalin-YFP.** z-stacks of 0.9 µm confocal sections through the entire depth of the cell were acquired by LSM at 0.1 frames s^-1^. Only the YFP-channel is shown. Label, min:s. Supports Fig. 7.

**Supplemental Movie 13. Actin dynamics in a non-growing pollen tube coexpressing NtPIP5K6-mCherry and mTalin-YFP.** z-stacks of 0.9 µm confocal sections through the entire depth of the cell were acquired by LSM at 0.1 frames s^-1^. Only the YFP-channel is shown. Label, min:s. Supports Fig. 7.

